# Rad55-Rad57 and Srs2 regulate homology search onset, coordination, reach and inactivation

**DOI:** 10.64898/2026.09.15.751666

**Authors:** Nicolas Mendiboure, Agnès Dumont, Jérôme Savocco, Chloé Dupont, Jie Liu, Daniel Jost, Wolf-Dietrich Heyer, Aurèle Piazza

## Abstract

DNA double-strand break (DSB) repair by homologous recombination entails the coordinated search for a homologous dsDNA molecule by two heterotypic Rad51-ssDNA filaments in eukaryotes. How homology search is regulated in cells remains largely unknown. Using genomic and molecular assays to track spatial chromatin organization and early recombination intermediates, we investigated the roles in homology search of two antagonistic regulators of Rad51-ssDNA filaments metabolism in *S. cerevisiae*: the Rad51 paralogs Rad55-Rad57 and the 3’-5’ ssDNA translocase Srs2. Srs2 promoted the coordinated search between filaments on each DSB ends, inactivated homology search following homology identification, and stimulated the first strand synthesis of BIR. Rad55-Rad57 both stimulated the formation of Rad51-ssDNA filaments and protected them against disruption by Srs2. Together, Rad55-Rad57 and Srs2 enacted a structural proof-reading that resulted in stiffer Rad51-ssDNA filaments competent for genome-wide homology search. This work reveals multiple ways by which the control of Rad51-ssDNA filament metastability by the individual and joint activities of Rad55-Rad57 and Srs2 regulate homology search onset, coordination, reach and inactivation.

## Introduction

Homologous recombination (HR) is a conserved DNA double-strand break (DSB) repair pathway that uses a double-stranded DNA (dsDNA) molecule of similar (*i*.*e*. homologous) sequence as a template. This intact information “donor” has to be identified in the nuclear space. In eukaryotes, the search for homology is catalyzed by heterotypic Rad51 filaments assembled on the ssDNA continuously generated by resection on each side of the DSB. Being part of the chromatin polymer, the exploratory capacity of these Rad51-ssDNA filaments is initially constrained to the immediate neighborhood of the DSB and influenced in *cis* by factors that organize chromosomes (Agmon *et al*, 2013; Renkawitz *et al*, 2013; Lee *et al*, 2016; Piazza *et al*, 2021; Marin-Gonzalez *et al*, 2025; Teloni *et al*, 2025; Piazza & Taddei, 2026; Piveteau *et al*, 2026). If no suitable donor is identified during this first local phase, homology search progressively expands to more distant chromosomal regions. Genome-wide mapping of ssDNA contacts coupled to polymer modelling indicated that the rigidification of ssDNA by Rad51 underlay the transition from local to genome-wide homology search (Dumont *et al*, 2024). This rigidification coincided with the cytological appearance of dynamic, micrometer-long Rad51 filament structures spanning across the budding yeast nucleus (Liu *et al*, 2023). Accordingly, Rad51 mutant proteins forming short filaments *in vitro* are defective for inter-homolog but not inter-sister recombination (Meyer *et al*, 2026), and mutants defective for long-range resection that fail to assemble long Rad51-ssDNA filaments also fail to engage distant genomic loci and are specifically defective in inter-chromosomal repair (Kimble *et al*, 2023; Liu *et al*, 2023; Dumont *et al*, 2024). Long, stiff Rad51-ssDNA filaments thus fulfill a main requirement for genome-wide homology search in the *in vivo* context in which chromatin loci are confined to small nuclear areas: by projecting DSB-proximal filament regions at a distance from the chromatin environment where the DSB occurred (Piazza & Taddei, 2026). These observations tie the competence for genome-wide homology search to the length and stiffness of Rad51-ssDNA filaments.

The high bending stiffness of RecA/Rad51-ssDNA filaments has long been recognized *in vitro* (Sheridan *et al*, 2008; Miné *et al*, 2007; Hegner *et al*, 1999; van der Heijden *et al*, 2007). *E. coli* RecA nucleates infrequently on ssDNA, but subsequent oligomerization is highly cooperative, which results in continuous filaments exhibiting high persistence length (Cazenave *et al*, 1983; Hegner *et al*, 1999; Galletto *et al*, 2006; Joo *et al*, 2006; Bell *et al*, 2012). Differently, yeast and human Rad51 nucleate more frequently on both ssDNA and dsDNA, but growth is comparatively slow, which results in a high degree of filament segmentation and overall reduced persistence length compared to RecA (Heuser & Griffith, 1989; Ogawa *et al*, 1993; Sung & Robberson, 1995; Mameren *et al*, 2006; Miné *et al*, 2007; Modesti *et al*, 2007; van der Heijden *et al*, 2007; Sheridan *et al*, 2008; Hilario *et al*, 2009; Liu *et al*, 2011; Muhammad *et al*, 2024). Ancillary factors that impinge on the intrinsic Rad51 nucleation, extension and dissociation rates on ssDNA are likely to modulate the degree of filament segmentation, its overall bending rigidity, and may thus regulate the competence for genome-wide homology search.

In *S. cerevisiae*, the metabolism of Rad51-ssDNA filaments is controlled by opposing activities imparted by the Rad51 paralogs Rad55-Rad57 on the one hand and the 3’-5’ motor protein and helicase Srs2 on the other hand (reviewed in (Bonilla *et al*, 2020)). Rad55-Rad57 and their homologs in *C. elegans* and *H. sapiens* associate to the 5’ side of Rad51 filaments, and have been reported to promote their formation on protein-free and RPA-coated ssDNA *in vitro* (Sung, 1997; Gaines *et al*, 2015; Roy *et al*, 2021; Belan *et al*, 2021; Greenhough *et al*, 2023; Akita *et al*, 2024; Deveryshetty *et al*, 2025; Greenhough *et al*, 2026; Koo *et al*, 2026; Rawal *et al*, 2026). This function may involve stimulating filament nucleation, stabilizing nuclei prior to their elongation, and/or promoting filament growth. Oppositely, Srs2 disrupts Rad51-ssDNA filaments in an ATPase-dependent manner (Veaute *et al*, 2003; Krejci *et al*, 2003, 2004; Le Breton *et al*, 2008; Antony *et al*, 2009; Kaniecki *et al*, 2017). This disruption reaction is counteracted by Rad52 and Rad55-Rad57 (Roy *et al*, 2021; Liu *et al*, 2011; Ma *et al*, 2018, 2021), which mechanistically may involve both (i) direct protection through their stable association with the Rad51-ssDNA filament and (ii) kinetic competition by promoting re-assembly of the filament (Hays *et al*, 1995; Fortin & Symington, 2002; Liu *et al*, 2011; Esta *et al*, 2013; Gibb *et al*, 2014; Ma *et al*, 2018; Roy *et al*, 2021; Belan *et al*, 2021; Maloisel *et al*, 2023). Consistent with these biochemical functions, deletion of *SRS2* suppressed the sensitivity of the *rad55*Δ and *rad57*Δ mutants to various genotoxic treatments that induce sister-based recombination, such as ionizing radiation (Fung *et al*, 2009; Liu *et al*, 2011; Xu *et al*, 2013; Elango *et al*, 2017). Differently, end-point assays specifically reporting on inter-homolog recombination revealed that defects of the *rad55*Δ and *rad57*Δ mutants were only partly suppressed by *SRS2* deletion (Elango *et al*, 2017; Maloisel *et al*, 2023). It suggested that cells defective for both Rad55-Rad57 and Srs2 exhibit a specific defect in achieving inter-chromosomal HR repair. The nature of this defect remains unknown, but could indicate a defective assembly of Rad51-ssDNA filaments competent for genome-wide homology search.

The two DNA ends that perform homology search remain tethered in space during HR, a phenomenon primarily studied in budding yeast (Kaye *et al*, 2004; Lobachev *et al*, 2004; Clerici *et al*, 2005; Nakai *et al*, 2011; Piazza *et al*, 2021; Phipps *et al*, 2024). End-tethering depends the catalytic activities of the short- and long-range resection factors Sae2-MRX and Exo1, respectively; an observation congruent between by both live microscopy tracking the distance between chromatin loci on each side of the DSB with fluorescent repressor-operator arrays and Hi-C (Kaye *et al*, 2004; Lobachev *et al*, 2004; Clerici *et al*, 2005; Nakai *et al*, 2011; Piazza *et al*, 2021; Phipps *et al*, 2024). The Ddc1-Mec3-Rad17 clamp (human 9-1-1 clamp) present at the ssDNA-dsDNA resection junction also promoted end-tethering, independently of its DNA damage checkpoint function (Piazza *et al*, 2021). These observations indicated that the dsDNA-ssDNA junction is a main tethering point between the two broken chromosomal fragments, from which Rad51-ssDNA filaments emanate. Accordingly, mapping of Rad51-ssDNA contacts revealed a partly coordinated search for homology between opposite DSB ends, which was abolished in the long-range resection- and end tethering-deficient *exo1* mutant (Dumont *et al*, 2024). Furthermore, cytological analysis of fluorescently labeled Rad51 predominantly formed a unique filament structure despite the presence of four DSB ends generated at a single site on replicated sister chromatids, which suggested that filaments were aligned during homology search (Liu *et al*, 2023). These observations raised the possibility that a function for DSB end-tethering could be to organize Rad51-ssDNA filaments for coordinated homology search. This suggestion remains to be established in a resection-proficient context. More broadly, it remains unclear whether Rad51-associated factors contribute to coordinate homology search by opposite filaments.

Homology search is conducted in parallel at multiple sites along RecA/Rad51-ssDNA filaments (Forget & Kowalczykowski, 2012). Homology identification only involves a sub-section of the ssDNA partaking in the search, resulting in ∼200 bp-long D-loops even on topologically unconstrained donor molecules, as well as multi-invasions resulting from independent homology identification events on distinct dsDNA molecules *in vitro* (Wright & Heyer, 2014; Piazza *et al*, 2017; Shah *et al*, 2020). Whether the Rad51-ssDNA filament sections outside the D-loop keep on searching for homology in cells, or whether a feedback mechanism exists that inactivates homology search upon homology identification, remains unknown.

Here we investigated the roles of Rad55-Rad57 and Srs2 in regulating homology search following site-specific DSB induction by tracking core HR intermediates and steps in *S. cerevisiae* cells: resection, Rad51-ssDNA filament formation, homology search, D-loop formation and D-loop extension. It confirmed prior suggestions that Rad55-Rad57 promotes the formation of Rad51 filaments and antagonizes Srs2 disruption activity in cells. It further revealed a role of Srs2 in promoting coordinated search between both DSB ends and in inactivating homology search once homology had been identified. The combined presence of Rad55-Rad57 and Srs2 promoted homology search at distant genomic sites, suggesting that they promote the formation of stiff Rad51 filaments. Accordingly, purified Rad55-Rad57 and Srs2 together reduced Rad51-ssDNA filaments segmentation, resulting in long and continuous filaments *in vitro*. Stochastic modeling suggests that such disruption-driven filament growth can be achieved if Rad55-Rad57 restricts the disruption activity of Srs2 to single Rad51-ssDNA segments, allowing for the growth of the nearby segments. These results reveal multiple ways by which Rad55-Rad57 and Srs2 individually and collectively control homology search: its enactment, the coordination of search between opposite filaments, the pace of the transition from local to genome-wide homology search, and search inactivation upon homology identification.

## Results

### Experimental system

Successful homology identification by the Rad51-ssDNA filament leads to the formation of a displacement loop (hereafter D-loop), a DNA joint molecule produced upon DNA strand invasion (Wright *et al*, 2018). The absolute amount of D-loops in the cell population and their extension by a DNA polymerase can be quantified following acute site-specific DSB induction using the D-loop-Capture (DLC) and D-loop extension (DLE) assays, respectively (**Fig. S1A**) (Piazza *et al*, 2018, 2019; Reitz *et al*, 2022). We employed a well-established site-specific DSB induction system in haploid *S. cerevisiae* cells, which yields >90% of broken DNA molecules at a HO cut-site introduced on chr. V within 1 hour of *HO* over-expression (Piazza *et al*, 2018, 2019). The left DSB end shares 2 kb of homology with two competing donors: an intra-chromosomal donor located in close proximity (∼80 kb) and on the same chromosomal fragment as the homology-containing left DSB end on chr. V; and an inter-chromosomal donor on chr. II (hereafter intra and inter donors, respectively; **Fig. 1A**) (Piazza *et al*, 2021; Djeghmoum & Piazza, 2025). This experimental system authorizes all HR steps up to, and including, D-loop extension. Downstream repair steps are purposefully precluded to prevent cell growth resumption: a lack of homology to the second end prevents regular gene conversion, and the orientation of the donor towards the centromere or the DSB itself prevents telomere capture and thus BIR completion (Pham *et al*, 2021; Morrow *et al*, 1997).

**Figure 1:**
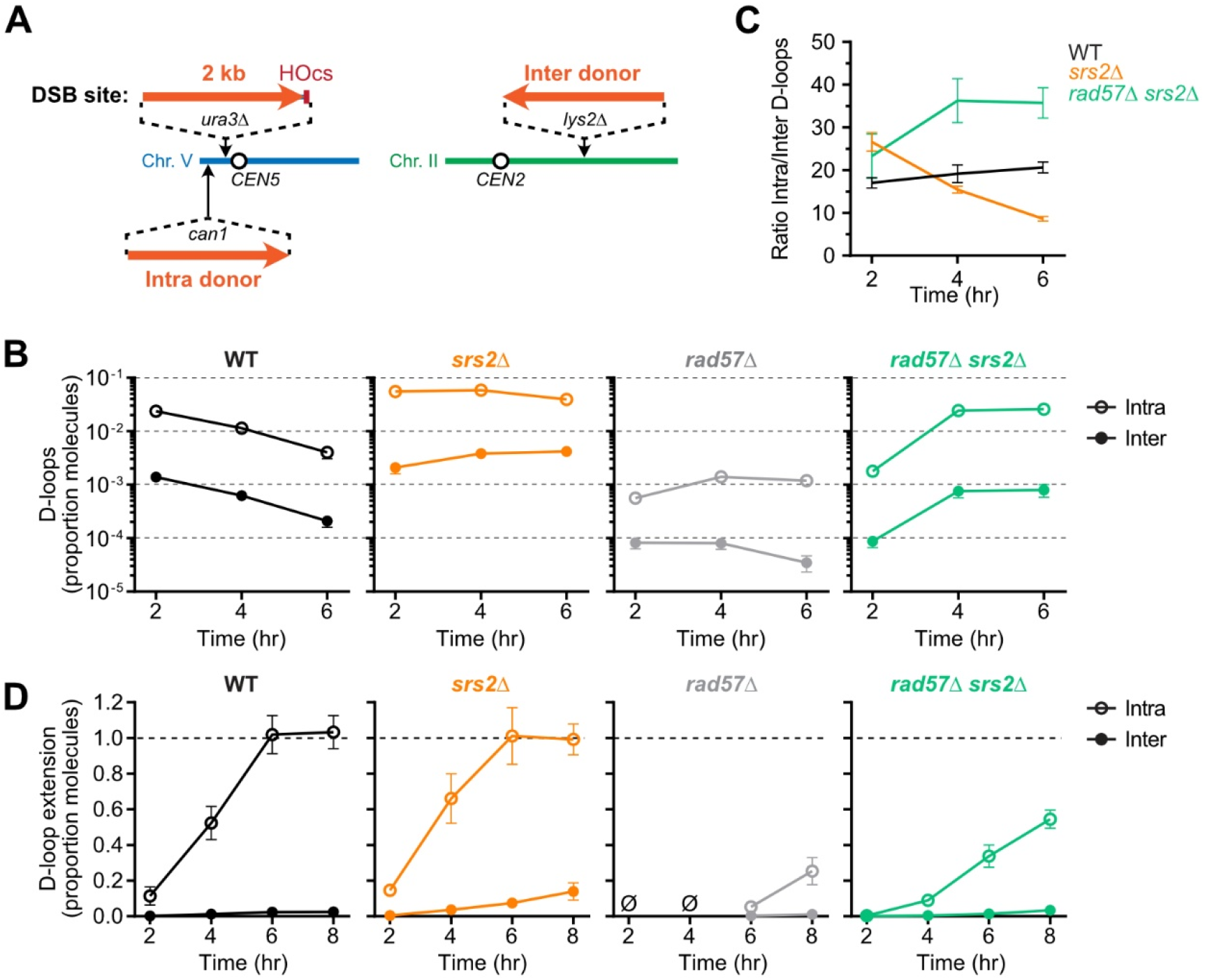
DSB-donor proximity partly alleviates the D-loop formation and extension defects of the *rad57*Δ *srs2*Δ mutant. (A) Unrepairable site-specific DSB induction system with two competing intra- and inter-chromosomal donors to the left DSB end. (B) Absolute D-loop levels at the intra- and inter-chromosomal donors in a WT (APY809), *srs2*Δ (APY1769), *rad57*Δ (APY1770), and *rad57*Δ *srs2*Δ (APY1772) strains. Data show mean ± SEM of at least 3 biological replicates. (C) Ratio of D-loops formed at the intra-over the inter-chromosomal donor, calculated from data in B). Data show mean ± SEM. (D) Absolute D-loop extension levels at the intra- and inter-chromosomal donors in the same strains as in B). Ø: no DLE products were detected at 2 and 4 hours in the *rad57*Δ mutant. Other legends as in B).

In the absence of break, the Hi-C contact frequencies determined between the DSB-inducible site and the intra donor is 13.4±1.1-fold higher than with the inter donor in asynchronous cells, and 29.7±7.7-fold higher in metaphase-arrested cells (*i*.*e*. the physiological context in which cells arrest and attempt HR repair following activation of the DNA damage checkpoint; **Fig. S1B**) (Piazza *et al*, 2021). Identification of the inter donor thus requires a broader exploration of the nuclear space than the identification of the intra donor. Accordingly, D-loops formed at the intra donor with faster kinetics and with a ∼20-fold preference relative to the inter donor in WT cells at all time points (**Fig. 1B-C** and **Fig. S1C**) (Piazza *et al*, 2021; Djeghmoum & Piazza, 2025), in general agreement with the Hi-C contact frequencies. This competitive system thus allows to distinguish general recombination defects, which should equally affect D-loop formation at both donors, and specific defects in genome-wide homology search, which should disproportionately inhibit D-loops at the inter donor. We used this system to gain insights into the role of Rad55-Rad57 and Srs2 in regulating homology search.

### Srs2 inactivates homology search following donor identification

D-loop levels were maximal at 2 hours post-DSB induction and decrease 10-fold within the next 4 hours in WT cells (**Fig. 1B**) (Piazza *et al*, 2021; Djeghmoum & Piazza, 2025). This decrease coincided with, and were previously shown to depend on, D-loop extension by PCNA/Pol*δ*, which was maximal at 6 hours post-DSB induction (**Fig. 1D**) (Reitz *et al*, 2023). ssDNA was not degraded and extension products were not converted to dsDNA in this system (**Fig. S1D**). Hence, despite the prolonged presence of ssDNA at all time points, D-loop levels were not maintained over time. It suggested that the competence of ssDNA for D-loop formation was lost following D-loop extension.

In contrast to WT cells, D-loop levels increased (inter) or remained constant (intra) in *srs2*Δ cells, involving up to 0.4 and 5.9% of broken molecules 6 hours post-DSB induction, respectively (**Fig. 1B**). This persistence of D-loops occurred in spite of the efficient D-loop extension taking place between 2 and 6 hours (**Fig. 1D**). These results indicate that Srs2 is required to abrogate the competence of ssDNA for invasion once D-loop formation or extension has occurred.

Additionally, Srs2-defective cells exhibited an exacerbated preference for the intra-over the inter-chromosomal donor at the earliest time point (26.6±3.7-fold, vs. 17.0±3.3-fold in WT cells; **Fig. 1C**). This preference progressively decreased over time, from 26.6±3.7-fold at 2 hours to 8.6±1.2-fold at 6 hours, significantly lower than the 20.6±3.9-fold preference observed in WT cells (p<0.001 two-tailed Welch’s t-test, **Fig. 1C**). These observations suggest that Srs2-deficient cells are initially defective for, but over time become unrestricted in long-range homology search.

We sought to determine the mechanism(s) by which Srs2 (i) abrogates ssDNA competence for D-loop formation following D-loop extension and (ii) prevents the erosion of the intra/inter donor preference over time.

### Rad55-Rad57 promotes homology search partly by counteracting Srs2

D-loops were reduced 20-to 40-fold in a *rad57*Δ mutant relative to WT cells at 2 hours post-DSB induction (**Fig. 1B**). Intra D-loops never exceeded 0.2% of total DNA molecules, while inter D-loops hovered close to the detection limit (∼10^-5^) (Reitz *et al*, 2023), which prevented a confident determination of the intra/inter donor preference in this mutant. Consistently, D-loop extension at the intra donor did not appreciably occur before 6 hours post-DSB induction, a >4 hours delay relative to WT cells (**Fig. 1D**). D-loop joint molecules and D-loop extension at both donors were similarly decreased relative to WT cells (**Fig. S1E-G**). A similar defect was observed in both the *rad55*Δ and *rad57*Δ mutants in an independent strain bearing a single inter-chromosomal donor (**Fig S2A-F**). Consequently, cells lacking Rad55-Rad57 are defective for donor identification irrespective of its location, indicative of a general recombination impairment.

This general defect was partly suppressed upon *SRS2* deletion: D-loop levels increased by an order of magnitude at 4 and 6 hours post-DSB induction (**Fig. 1B**), and extension products were detected at both donors (**Fig. 1D**) in the *rad57*Δ *srs2*Δ mutant, reaching 56.9 ± 10.9% of molecules at 8 hours post-DSB induction. This partial recovery was nonetheless delayed both for D-loop joint molecule formation and extension relative to WT and *srs2*Δ strains (**Fig. 1B, D**). This D-loop formation and extension delay was confirmed in both *rad55*Δ *srs2*Δ and *rad57*Δ *srs2*Δ strains bearing a single inter-chromosomal donor (**Fig. S2A-F**).

These observations confirm (i) the role of Rad55-Rad57 in counteracting the anti-recombination activity of Srs2 and (ii) suggest an additional function of Rad55-Rad57, or of its interplay with Srs2, in accelerating the kinetics of D-loop formation.

### Proximity with the donor partly alleviated D-loop formation defects in the *rad57*Δ *srs2*Δ mutant

Interestingly, the bias toward the intra donor was exacerbated in the *rad57*Δ *srs2*Δ mutant relative to WT or *srs2*Δ cells (*p*<0.05 at both 4 and 6 hours post-DSB induction, two-tailed Welch’s t-test, **Fig. 1B-C**). Likewise, D-loop formation and extension defects were less pronounced at the intra than at the inter donor in the *rad57*Δ *srs2*Δ mutant relative to WT or *srs2*Δ cells (**Fig. S1E-G**). The spatial proximity of the donor thus partly alleviated a recombination defect specific to the *rad57*Δ *srs2*Δ mutant. Hence, in addition to a general delay in homology search onset, *rad57*Δ *srs2*Δ cells exhibited a specific defect in identifying spatially distant homologies.

In the following sections, we sought to determine the upstream molecular basis of both the general and specific HR defects determined at the D-loop level in the *rad57*Δ, *srs2*Δ, and *rad57*Δ *srs2*Δ mutants by quantifying the presence of Rad51 on dsDNA and ssDNA by strand-specific calibrated ChIP-seq, the activity and contacts made by these filaments with ssHi-C, and the overall chromosome organization with Hi-C. We note that none of these mutants have notable resection defects (see below, **Fig. 2B, S4B** and **Supplementary Code 1**).

**Figure 2:**
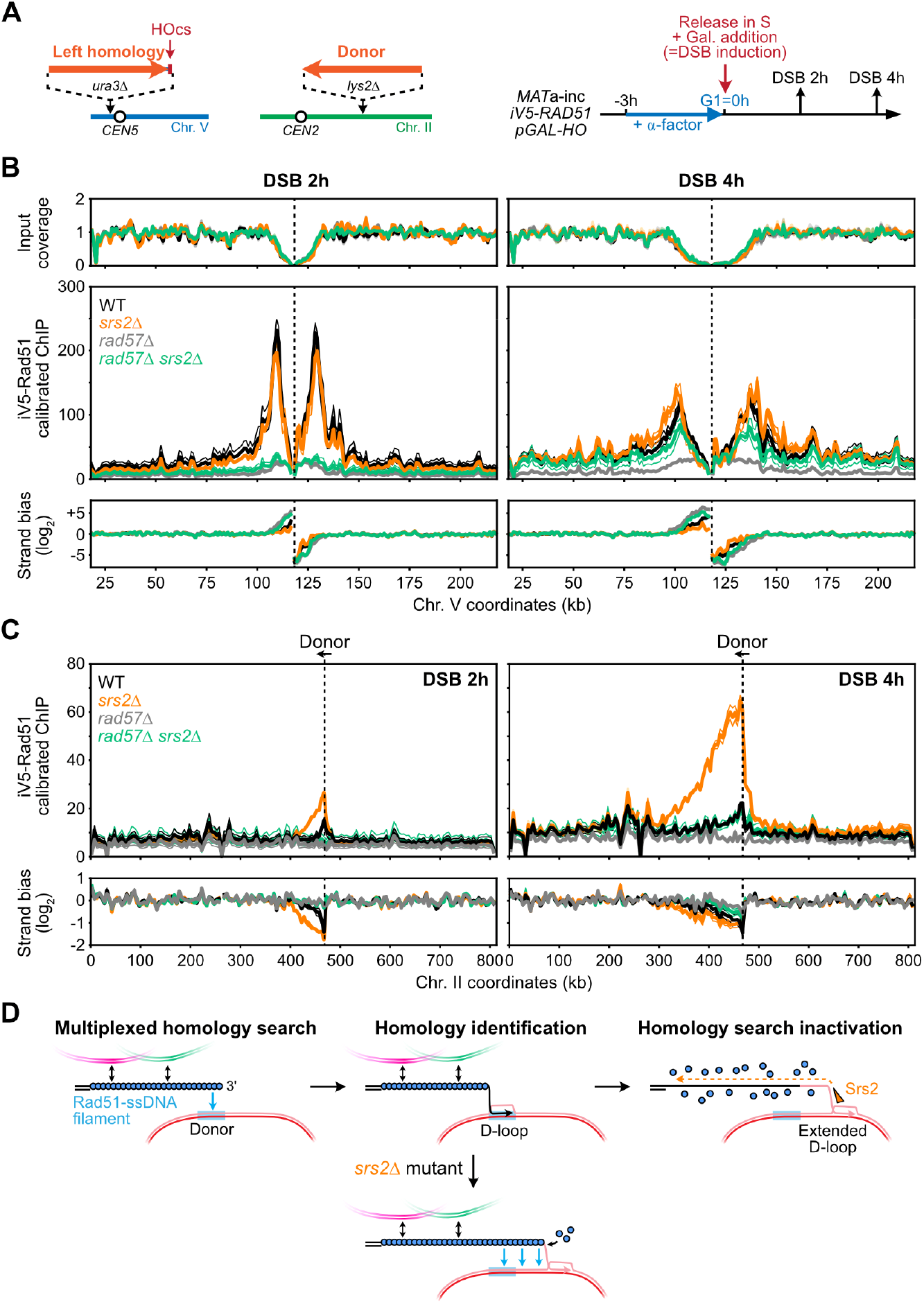
Rad55-Rad57 and Srs2 control the formation and the post-synaptic disassembly of Rad51-ssDNA filaments. (A) DSB-inducible system and synchronization scheme. Only the left DSB end has homology to a donor on chr. II. (B) Rad51 is recruited on ssDNA on each side of the DSB. Strand-specific calibrated ChIP-Seq of iV5-Rad51 in the 200 kb region surrounding the DSB site in WT (APY1895), *srs2*Δ (APY2033), *rad57*Δ (APY2029), and *rad57*Δ *srs2*Δ (APY2037). The strand bias reveals the binding of Rad51 on resected ssDNA. Data show mean ± SEM of n=3 (WT) and n=2 (other genotypes) biological replicates. Data are binned at 1 kb. (C) Asymmetric recruitment of Rad51 in the chromosomal region downstream of the donor site on chr. II, from data in B). Data are binned at 5 kb. The average value per kb is plotted. (D) Model for Srs2-mediated inactivation of homology search upon Rad51 removal from the ssDNA upstream of the extended D-loop.

### The formation of Rad51 filaments is delayed in the *rad57*Δ *srs2*Δ mutant

The dysregulated D-loop levels in *rad57*Δ and *srs2*Δ mutants may originate from defects in Rad51-ssDNA filaments formation/stability and elimination, respectively. Rad51 filaments in WT cells and these mutants have only be monitored by semi-quantitative and/or non-calibrated chromatin immunoprecipitation (ChIP) (Sugawara *et al*, 2003; Renkawitz *et al*, 2013; Peritore *et al*, 2021), or indirectly by measuring the formation of Rad51 foci in cells (Gasior *et al*, 2001; Lisby *et al*, 2004; Fung *et al*, 2006). To overcome these limitations, we carried out strand-specific calibrated ChIP-seq against Rad51 bearing an internal V5 epitope at 2 and 4 hours post-DSB induction. The V5 epitope was introduced internally at a safe-harbor site previously used to achieve functional tagging of Rad51 with GFP, between the G54 and G55 residues in the N-terminal tail of Rad51 at its endogenous locus (iV5-Rad51; see **Methods**) (Liu *et al*, 2023; Deveryshetty *et al*, 2025). Resistance to camptothecin confirmed that this iV5-Rad51 construct was largely functional (**Fig. S3A**). *C. glabrata* bearing a V5-tagged Scc1 protein was used as a calibrator (Hu *et al*, 2015).

DSB formation was induced upon S-phase release from a G1 block, yielding highly homogeneous populations of metaphase-arrested cells undergoing HR repair 2 and 4 hours later in all strains (**Fig. 2A** and **S3B**) (Dumont *et al*, 2024). Rad51 was strongly enriched within a ∼30 kb region on each side of the DSB in WT cells (**Fig. 2B**). A strong strand bias matching the position, orientation and length of the resection tracts indicated that at least a fraction of the ChIP signal originated from Rad51 binding to ssDNA (**Fig. 2B**). Rad51 recruitment in the *rad57*Δ mutant was reduced ∼5-fold relative to WT cells and remained confined to the immediate vicinity of the DSB (**Fig. 2B** and **S3C**). Absence of Srs2 modestly increased Rad51 binding in the vicinity of the DSB at 4 hours post-DSB induction (**Fig. 2B**). Rad51 recruitment at the DSB in the *rad57*Δ *srs2*Δ mutant resembled that of the *rad57*Δ mutant at 2 hours, and increased to reach levels close to that observed in WT and *srs2*Δ strains at 4 hours post-DSB induction (**Fig. 2B**). These results are consistent with Rad55-Rad57 both (i) promoting formation of Rad51-ssDNA filaments and (ii) antagonizing their disruption by Srs2.

### Srs2 clears Rad51 from neo-synthesized ssDNA

Previous work revealed persistent DNA joint molecules formed during BIR in Srs2-deficient cells (Elango *et al*, 2017). Accordingly, D-loop levels at the donor site remained elevated in the *srs2*Δ mutant even after D-loop extension had occurred, unlike in a WT cells (**Fig. 1B, C**). These observations pointed at a post-D-loop extension defect: either a failure to prevent assembly of Rad51 filaments from extended ssDNA, and/or a failure to turnover DNA joint molecules resulting from the activity of these filaments (Elango *et al*, 2017; Liu *et al*, 2017).

ChIP-Seq of iV5-Rad51 revealed a broad enrichment region on chr. II that stretched asymmetrically from the donor with the directionality expected from recombination-associated DNA synthesis initiated at that site in WT and *srs2*Δ cells (**Fig. 2C**). Strand bias indicated that the majority of Rad51 was associated to the neo-synthesized DNA generated by BIR (**Fig. 2C**). Srs2 suppressed Rad51 binding up to 5-fold over the entire BIR track relative to WT cells (**Fig. 2C**). The Srs2-mediated suppression of Rad51 binding to the extended broken molecule likely underlies the suppression of D-loops formed post-D-loop extension in WT cells (**Fig. 2D**).

#### Srs2 inhibits the rate of D-loop extension during BIR

The Rad51-bound region downstream of the donor increased from ∼50 and ∼140 kb from 2 to 4 hours post-DSB induction in WT cells and from ∼70 to ∼200 kb in *srs2*Δ cells (**Fig. 2C**). These measurements yielded a maximum leading-strand BIR synthesis rate of ∼45 kb/hour in WT cells and ∼65 kb/hour in the *srs2*Δ mutant. The WT rate is in general agreement with the average BIR rate (∼30 kb/hour) previously deduced by quantifying the DNA copy number gain at increasing distances from the invasion point over time (Liu *et al*, 2021). The ∼20 kb/hr increase observed in the *srs2*Δ mutant indicates that Srs2 inhibits the first strand synthesis of BIR; an activity consistent with its ability to dissociate extended D-loops *in vitro* and in cells (Liu *et al*, 2017; Piazza *et al*, 2019; Hung *et al*, 2025).

### General chromosome organization is unaffected in the *rad57*Δ, *srs2*Δ and *rad57*Δ *srs2*Δ mutants

We next sought to investigate the nature of (i) the progressive loss of intra-chromosomal donor preference in the absence of Srs2 and (ii) the specific defect in inter-chromosomal homology search in the combined absence of Rad55-Rad57 and Srs2. Since chromosome organization impinges on the search process (Agmon *et al*, 2013; Lee *et al*, 2016; Piazza *et al*, 2021; Dumont *et al*, 2024; Piazza & Taddei, 2026), we first determined whether the lack of Srs2 and/or of Rad55-Rad57 impacted chromosome organization by performing Hi-C at 2 and 4 hours post-DSB induction in the relevant mutants. DSB formation was induced in a cell population synchronously released from G1, as for ChIP-seq (**Fig. 2A**), leading to a homogeneous population of G2/M-arrested cells 2 and 4 hours later (**Fig. S4A**). None of the mutants assayed (*rad51*Δ, *rad57*Δ, *srs2*Δ, and a combination thereof) caused notable changes to DNA damage-induced G2/M-arrest, resection tract length, genome-wide cohesin-mediated chromatin loop folding and centromere clustering (a main feature of the Rabl chromosome organization in yeast; **Fig. S4A-F**). Consequently, dysregulated intra/inter donor identification in the *srs2*Δ and *rad57*Δ *srs2*Δ mutants (**Fig. 1C**) does not originate from a general alteration of chromosome organization.

### Srs2 promotes DSB end-tethering

Intriguingly, Hi-C revealed reduced DSB end-tethering in the *srs2*Δ mutant (**Fig. 3A**). This effect was not restrained to the immediate vicinity of the DSB, but involved the entire left and right chromosomal fragments, whose contact frequency significantly decreased from ∼30% at 2 hours to ∼70% at 4 hours relative to WT cells (**Fig. 3A** and **S5A, B**). Consistently, the left, centromere-devoid chromosomal fragment lost its preferential interaction with the centromere of other chromosomes (**Fig. 3B**). The ATP hydrolysis-deficient *srs2-K41A* mutant (Krejci *et al*, 2004) resembled the *srs2*Δ mutant (**Fig. 3A, B** and **S5B**), suggesting that the translocase activity of Srs2 is required for its function in DSB end-tethering. The reduction in end-tethering in *srs2*Δ cells depended on Rad51 and Rad55-Rad57 (**Fig. 3C, D** and **S5C**). However, it did not depend on the presence of a donor (**Fig. S5D, E**) or of Pol3/Rfc1 (**Fig. S6**), which ruled out synaptic and post-synaptic roles of Srs2 in promoting end-tethering.

**Figure 3:**
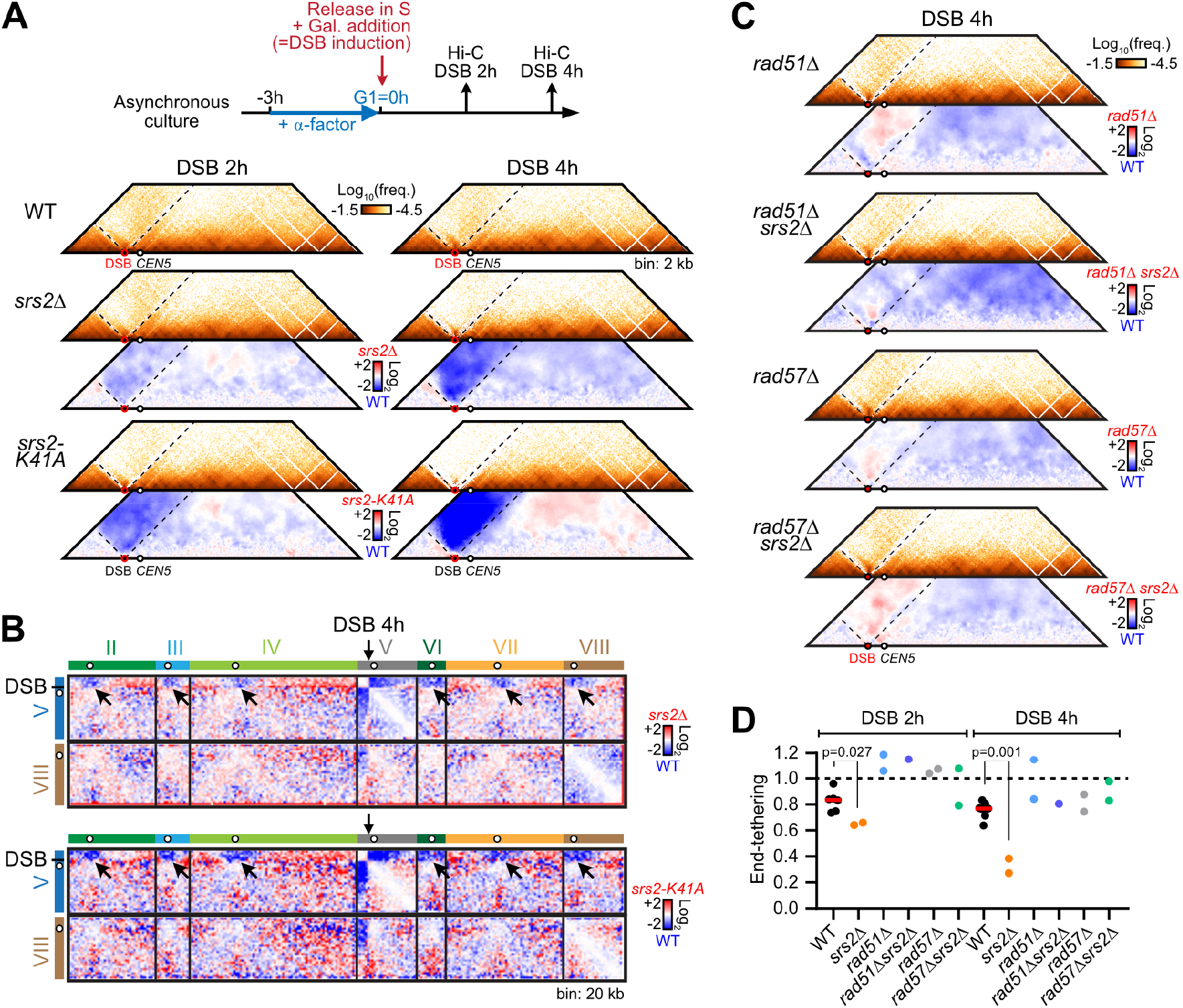
Loss of DSB end-tethering in Srs2-deficient cells. (A) Hi-C maps of chr. V in WT (APY266, n=6 or 7), *srs2*Δ (APY773, n=2), *srs2-K41A* (WDHY4616, n=1) strains, from data subsampled to 24 million contacts and binned at 2 kb. Ratio maps over WT are shown, which highlights the loss of contacts between the left and right chromosomal fragments. (B) Ratio maps of intra- and inter-chromosomal Hi-C contacts in *srs2*Δ and *srs2-K41A* mutants over a WT strain highlighting the loss of contacts between the left, centromere-devoid chr. V fragment with other centromeres. The intact chr. VIII is shown for comparison. Same data as in A). (C) Hi-C maps of chr. V in the *rad51*Δ (APY679, n=2), *rad51*Δ *srs2*Δ (APY1666, n=1), *rad57*Δ (APY654, n=2), *rad57*Δ *srs2*Δ (APY1409, n=2) mutants and corresponding ratio maps over a WT strain (APY266) at 4 hours post-DSB induction. (D) Quantification of DSB end-tethering in WT (APY266), *srs2*Δ (APY773), *rad51*Δ (APY679), *rad51*Δ *srs2*Δ (APY1666), *rad57*Δ (APY654), and *rad57*Δ *srs2*Δ (APY1409) strains as described in **Fig. S5A**. Data points show individual biological replicates. Bar shows the median. Distributions were compared using an unpaired two-tailed Student t-test without Welch’s correction.

Cohesin is enriched in the DSB-surrounding chromatin region and was shown to regulate the initial homology search phase (Ström *et al*, 2004; Ünal *et al*, 2004; Dauban *et al*, 2020; Piazza *et al*, 2021; Scherzer *et al*, 2022; Dumont *et al*, 2024). We addressed its involvement in end-tethering upon auxin-induced depletion of its AID-tagged Scc1 subunit at the time of DSB induction (**Fig. S7A**). As expected, this depletion resulted in loss of chromatin loops between cohesin-associated regions (CARs) genome-wide in otherwise WT or mutant backgrounds without affecting the G2/M arrest (**Fig. S7B-D**) (Dauban *et al*, 2020; Costantino *et al*, 2020; Piazza *et al*, 2021). Cohesin loss did not affect end-tethering in an otherwise WT background (**Fig. S7E, F**). Absence of Srs2 caused a Rad51- and Rad57-dependent reduction of end-tethering in Scc1-depleted cells (**Fig. S7E, F**), as observed in cohesin-proficient cells (**Fig. 3D**). Cohesin thus does not play any appreciable role in DSB end-tethering, both in WT and *srs2*Δ cells.

These observations show that the ATPase activity of Srs2 is required to counteract a progressive loss of end-tethering originating in a pre-synaptic activity of the Rad51-ssDNA filament. This loss of end-tethering releases the left chromosomal fragment from its preferential association with other centromeres, granting access to more interstitial chromosomal regions (**Fig. 3B, G**). It is likely to contribute to the disproportionate increase of D-loops formed at the inter donor observed at 4 hours by DLC (**Fig. 1B, C**, and see below).

### Srs2 promotes coordinated search between DSB ends

To address the functional consequences of this loss of end-tethering, we monitored homology search using ssDNA-specific Hi-C (ssHi-C), a Hi-C-based method that retrieves contacts made at selected ssDNA sites (**Fig. 4A**) (Dumont *et al*, 2024). We determined the genome-wide contacts made by 20 ssDNA sites located on each side of, and up to 17.2 kb away from the DSB (**Fig. 4B**), as well as at 7 intact dsDNA sites on other chromosomes used as capture controls (Dumont *et al*, 2024) (**Methods**). Unlike Rad51 ChIP-Seq, which has often been used as a homology search readout (Renkawitz *et al*, 2013; Peritore *et al*, 2021; Teloni *et al*, 2025; Marin-Gonzalez *et al*, 2025), ssHi-C can resolve contacts made by the left or right Rad51-ssDNA filaments (**Fig. 4C**). The distribution of ssDNA contacts can be compared to that of intact dsDNA sites (**Fig. 4C**), allowing to infer some of its physical properties (see below) (Dumont *et al*, 2024).

**Figure 4:**
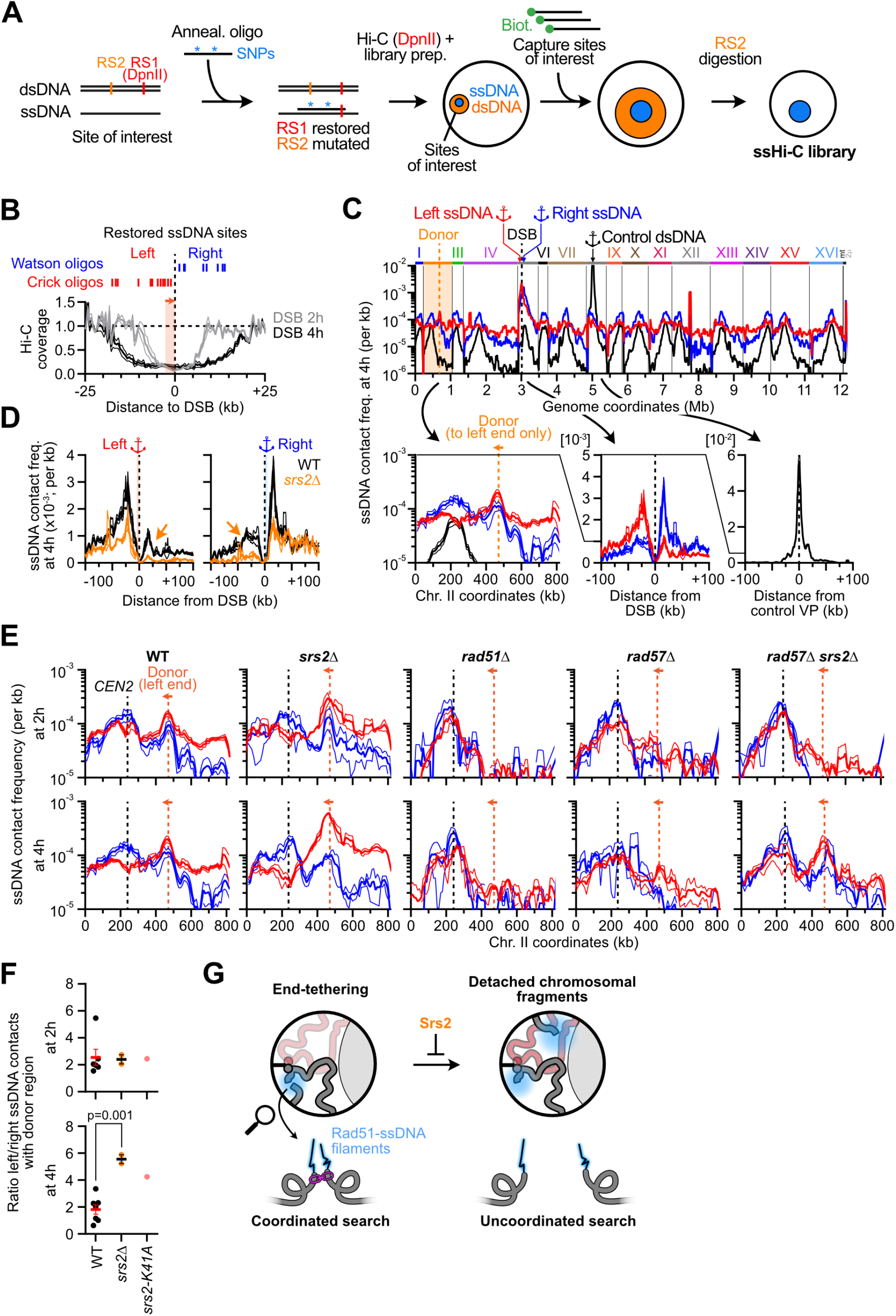
Srs2 promotes coordinated search between opposite Rad51-ssDNA filaments. (A) Overview of the ssHi-C procedure, starting from a crosslinked cell culture. RS1: DpnII. RS2: MfeI/SspI. ssDNA contacts are identified as reads bearing SNPs introduced in the annealing oligonucleotides. (B) Hi-C coverage in WT cells and position of the ssDNA sites restored on each side of the DSB. (C) Genome-wide 4C-like ssHi-C contact profiles of the average of the left ssDNA sites, right ssDNA sites, and of a control undamaged site 4 hours post-DSB induction in a WT strain (APY266). Data show mean ± SEM of n=7 biological replicates. Red: left ssDNA. Blue: right ssDNA. Black: control dsDNA site. (D) Distribution of left and right DSB-proximal ssDNA contacts in the 200 kb region surrounding the DSB site in a WT (APY266) and a *srs2*Δ (APY773) strain. Data show mean (± SEM) of n=7 and 2 biological replicates, respectively. Data are binned at 1 kb and smoothed over 5 bins. (E) Distribution of left and right DSB-proximal ssDNA contact along chr. II, which bears the donor to the left end of the DSB, in WT (APY266), *srs2*Δ (APY773), *rad51*Δ (APY679), *rad57*Δ (APY654) and *rad57*Δ *srs2*Δ (APY1409) strains. Data show mean ± SEM of n=2 biological replicates, except for WT (n=7). Data are binned at 10 kb and smoothed over 5 bins. (F) Ratio of the left and right ssDNA contacts with the donor region, from data in **Fig. S8C**. Data show individual biological replicates and the mean ± SEM. Distributions were compared using an unpaired two-tailed Student t-test without Welch’s correction. (G) Srs2 promotes DSB end-tethering and thus the coordination of homology search between both DSB ends. Loss of DSB end-tethering in Srs2-deficient cells causes a detachment of the centromere-devoid left chromosomal fragment from the rest of chr. V and its diffusion in the nucleoplasm.

ssDNA contacts were mainly enriched in the ∼200 kb DSB-surrounding region, with a side-specific preference: the left ssDNA contacted preferentially the left chromosomal fragment, and vice-versa (**Fig. 4C, D**). This bias was exacerbated in the *srs2*Δ mutant (**Fig. 4D**) and reduced in the *rad51*Δ and *rad57*Δ mutants (see below). These observations are in line with the HR-dependent reduction in end-tethering measured with Hi-C, and its exacerbation in Srs2-deficient cells (**Fig. 3A-D** and **S5B, C**).

ssDNA contacts were also enriched in a ∼100 kb region surrounding the donor site on chr. II for the left DSB end (**Fig. 4C, E**). These ssDNA-donor contacts increased from 2 to 4 hours and were homology, Rad51- and Rad57-dependent (**Fig. 4E** and **S8A-C**), confirming that they represent *bona fide* HR-dependent interactions (Dumont *et al*, 2024). These ssDNA-donor contacts were exacerbated in the *srs2*Δ mutant in a Rad51- and homology-dependent manner, and delayed in the *rad57*Δ *srs2*Δ mutant (**Fig. 4E** and **S8A-C**), consistent with DLC and Rad51 ChIP-seq data (**Figs. 1** and **2C**) (Piazza *et al*, 2019).

Notably, the homology-devoid right DSB end also engaged the donor (**Fig. 4C, E** and **S8A-C**), revealing a general coordination of homology search between both DSB ends, as noted previously (Dumont *et al*, 2024). This coordination is not absolute, as the right DSB end engaged the donor region ∼1.8±0.4 fold less than the homology-containing left DSB end in WT cells 4 hours post-DSB induction (**Fig. 4E, F** and **S8C**). This differential left/right interaction with the donor region was exacerbated ∼3-fold in the *srs2*Δ mutant 4 hours post-DSB induction (**Fig. 4F** and **S8C**). A similar decrease was observed in the *srs2-K41A* mutant (**Fig. 4F** and **Fig. S8A-C**). This decrease was commensurate with the end-tethering defect measured in these mutants at 4 hours (**Fig. 3** and **S5**). Consequently, Srs2 promotes coordinated search between the left and right Rad51-ssDNA filaments, and this coordination is correlated with the end-tethering determined by Hi-C (**Fig. 4G**).

Depletion of Scc1 did not affect search coordination, neither in a WT nor in a *srs2*Δ background (**Fig. S8D**), confirming that cohesin is dispensable for end-tethering and homology search coordination between opposite Rad51-ssDNA filaments (Piazza *et al*, 2021; Dumont *et al*, 2024).

### Rad55-Rad57 and Srs2 together promote distant homology search

Genome-wide profiling of ssDNA contacts on both DSB ends revealed a striking property of Rad51-ssDNA filaments: the capacity to frequently engage distant inter-chromosomal sites at the expanse of dsDNA regions flanking the resection tracts (**Fig. 4C, 5A** and **S9A**)(Dumont *et al*, 2024). The *rad57*Δ mutant was as defective as a *rad51*Δ mutant in engaging distant inter-chromosomal regions (**Fig. 5A** and **S9A**), consistent with a lack of Rad51-ssDNA filament in this mutant (**Fig. 2B**). Differently, the left and right ssDNA contact profiles were highly differentiated in the *srs2*Δ mutant 4 hours post-DSB induction: the left ssDNA engaged the rest of the genome uniformly while the right ssDNA exhibited a contact profile similar to that observed in WT cells (**Fig. 5B**). This differentiation is consistent with the loss of end-tethering and the uncoordinated search between both DSB ends observed in this mutant (**Fig. 3** and **4**), with the centromere-devoid left chromosomal fragment diffusing in the nucleoplasm (**Fig. 3G**). Finally, the genome-wide ssDNA contact profiles in the *rad57*Δ *srs2*Δ mutant resembled that of HR-deficient *rad51*Δ, *rad57*Δ and *rad51*Δ *srs2*Δ mutants at 2 hours post-DSB induction (**Fig. 5B**), consistent with the defect in Rad51 recruitment at the DSB observed by ChIP-seq (**Fig. 2B**). At 4 hours, inter-chromosomal contacts were marginally increased relative to HR-deficient cells (**Fig. 5B**), despite largely restored Rad51-ssDNA filaments at this time point (**Fig. 2B**).

**Figure 5:**
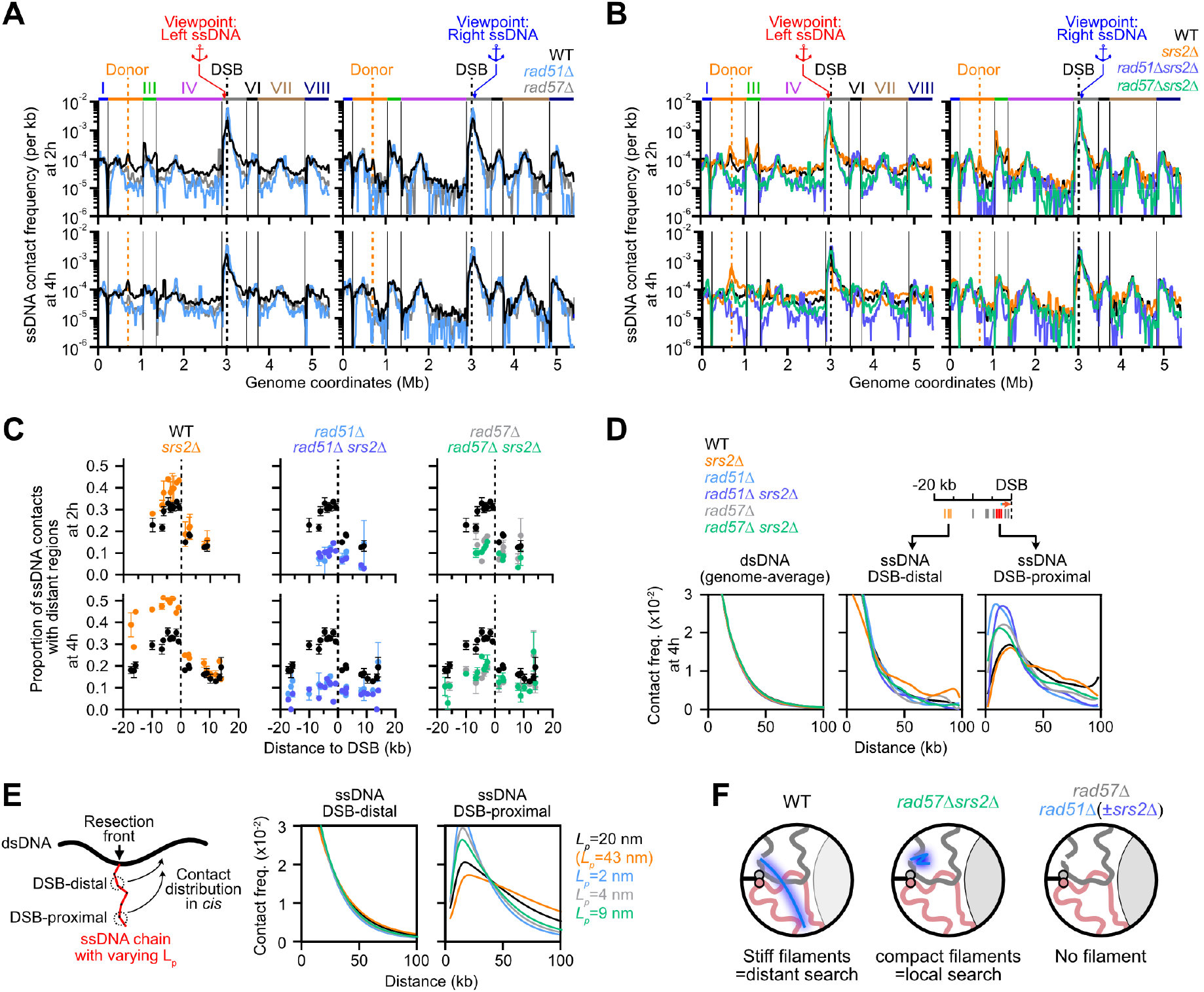
Rad55-Rad57 and Srs2 together promote genome-wide homology search. (A) 4C-like ssHi-C contact profiles of DSB-proximal left and right ssDNA sites in WT (APY266, n=7), *rad51*Δ (APY679, n=2) and *rad57*Δ (APY654, n=2) strains. Data show mean ± SEM. Data are binned at 10 kb and smoothed over 5 bins. (B) Same as A) in a *srs2*Δ (APY773, n=2), *rad51*Δ *srs2*Δ (APY1666, n=1) and *rad57*Δ *srs2*Δ (APY1409, n=2) strains. (C) Proportion of ssHi-C contacts with distant genomic regions for individual ssDNA sites in a WT, *srs2*Δ, *rad51*Δ, *rad51*Δ *srs2*Δ, *rad57*Δ and *rad57*Δ *srs2*Δ strains, from data in A) and B). Data show mean ± SEM. (D) Left: Contact frequency as a function of genomic distance for dsDNA genome-wide, DSB-distal ssDNA and DSB-proximal ssDNA on the left end side of the DSB in WT, *srs2*Δ, *rad51*Δ, *rad51*Δ *srs2*Δ, *rad57*Δ and *rad57*Δ *srs2*Δ strains 4 hours post-DSB induction, from data in A) and B). (E) Left: Rationale of the gaussian contact probability model. Right: Best-fit effective persistence length *L*_*p*_ for DSB-distal and DSB-proximal ssDNA data in D) (**Methods, Supplementary Code 1**). No *L*_*p*_ could fit both the DSB-distal and DSB-proximal ssDNA contact distributions obtained in the *srs2*Δ mutant. (F) Model for Rad51-ssDNA filament stiffness and exploratory capacity in WT and mutant cells.

ssHi-C can resolve the contacts made by individual sites along each ssDNA molecule, which we used to gain additional insights in the homology search defect of the *rad57*Δ *srs2*Δ mutant. In WT cells, DSB-proximal ssDNA sites engaged distant chromosomal regions more frequently than DSB-distal sites (**Fig. 5C**) at the expense of contacts with the flanking chromatin (**Fig. S9A**). Likewise, the distribution of contacts in *cis* spread at much longer range for DSB-proximal than for DSB-distal ssDNA sites, the latter of which exhibited a distribution similar to that of intact dsDNA sites (**Fig. 5D** and **S9B**). The DSB-proximal and DSB-distal ssDNA sites are thus contacting different chromosomal regions, which implies that they are spatially apart from each other on average in a WT cell population. Both the propensity for distant contacts and long-range *cis* contacts of DSB-proximal ssDNA sites depended on Rad51 and Rad57 (**Fig. 5C, D** and **S9B**), independently from the presence of a donor site (**Fig. S9C, D**). These observations implicated Rad51-ssDNA filaments in physically separating the DSB-proximal ssDNA sites from the resection front (Dumont *et al*, 2024).

Separation between the DSB-proximal and DSB-distal ssDNA sites can be achieved if ssDNA is stiffened by Rad51 relative to RPA-coated ssDNA, as is the case *in vitro*. Coupled to a stochastic description of resection, a Gaussian polymer model could infer the population-averaged effective persistence length (L_p_) of ssDNA by fitting the distribution of *cis* contacts for ssDNA sites on the left of the DSB as a function to their distance from the resection front (**Fig. 5E, Methods**) (Dumont *et al*, 2024). This yielded a best-fit L_p_ of 20 nm in WT cells at 4 hours post-DSB induction (**Fig. 5E** and **S9E**), consistent with our previous estimate (Dumont *et al*, 2024). In contrast, L_p_ was 2 nm in Rad51-deficient cells and 4 nm in the *rad57*Δ mutants. Contact distributions in the *rad57*Δ *srs2*Δ mutant yielded a L_p_ of 9 nm, only modestly increased relative to the *rad57*Δ mutant (**Fig. 5E** and **S9E**). It pointed at a substantial defect in ssDNA stiffening in the combined absence of Rad55-Rad57 and Srs2 despite the restoration of Rad51 recruitment at the DSB measured by chromatin immunoprecipitation (**Fig. 2B**). These results suggest that the combined presence of Rad55-Rad57 and Srs2 leads to a structural change in Rad51 filaments resulting in a stiffening of ssDNA, which supports genome-wide homology search (**Fig. 5F**).

Intriguingly, the *srs2*Δ data could not be satisfyingly fitted by the model, as no L_p_ value could satisfyingly account for the distribution of contacts made by both the DSB-distal and DSB-proximal ssDNA sites (**Fig. 5E** and **S9E**). It suggested that ssDNA stiffening was not uniform, being on average greater near the resection front than at other ssDNA regions. This issue is unlikely to result from the loss of end-tethering, as the model only considers the *cis* distribution of left ssDNA contacts with the left chromosomal fragment (see **Methods**).

### Rad55-Rad57 and Srs2 promote the assembly of continuous Rad51-ssDNA filaments *in vitro*

We sought to directly address the role of Rad55-Rad57 and Srs2 in altering the structural properties of Rad51-ssDNA filaments with purified proteins *in vitro*. Rad51 filaments assembled onto a 5.4 kb-long ssDNA molecule with ATP in the presence of absence of Rad55-Rad57 and/or Srs2 were imaged by negative-stained electron microscopy (**Fig. 6A**). Alone, Rad51 assembled into multiple segments interspersed with RPA-coated ssDNA regions that covered ∼17% of the ssDNA (471±185 nm-long segments on average vs. ∼ 2,700 nm assuming an ATP-bound Rad51-ssDNA contour length, **Fig. 6A-C**). Addition of Rad55-Rad57 did not increase the number of segments but decreased their average length (258±112 nm, p<0.0001 vs. Rad51 alone, Kolmogorov-Smirnov test; **Fig. 6A-C**). Differently, Srs2 disrupted most Rad51-ssDNA filaments in the absence of Rad55-Rad57 (**Fig. 6A-C**). The length of the surviving Rad51-ssDNA filaments was not significantly different from that observed in its absence (345±191 vs. 471±185 nm, **Fig. 6C**), suggesting that Srs2 eliminates all or most of the Rad51 segments from a ssDNA molecule at once (Kaniecki *et al*, 2017; Krejci *et al*, 2003; Veaute *et al*, 2003). As previously observed with shorter ssDNA substrates, Rad51 filaments assembled in the presence of Rad55-Rad57 were resistant to disruption by Srs2 (**Fig. 6A, B**) (Liu *et al*, 2011). The length of Rad51-ssDNA filaments assembled in the combined presence of Rad55-Rad57 and Srs2 reached 1343±785 nm, 3-fold longer than with Rad51 alone, and 5.2-fold longer than in the presence of Rad51 and Rad55-Rad57 (p<0.0001 in all cases, **Fig. 6A-C**). This increase was not driven by an increase in segment number (**Fig. 6B**), but by their individual length (**Fig. 6C**). Strikingly, Rad55-Rad57 and Srs2 led to the assembly of long (>500 nm) continuous Rad51-ssDNA segments, which accounted for the majority of the total filament length increase (**Fig. 6C**). This increase was not expected from the sole protection function for Rad55-Rad57, which should have resulted in a greater surviving fraction of Rad51 segments without a change to the segment length distribution. Likewise, it was not expected from a sole role in promoting Rad51 filament nucleation. It instead indicated a more dynamic assembly process enacted in the combined presence of Rad55-Rad57 and Srs2.

**Figure 6:**
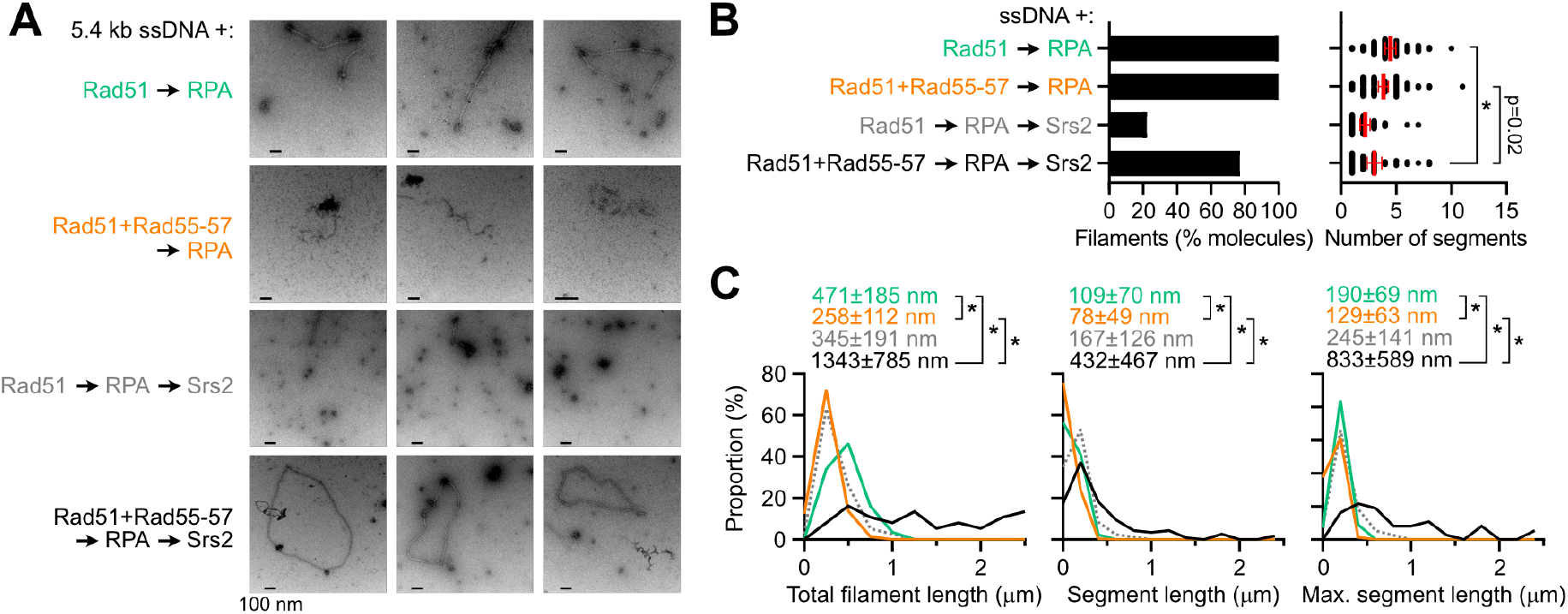
Rad55-Rad57 and Srs2 promote assembly of continuous Rad51-ssDNA filaments *in vitro*. (A) Representative electron micrographs of 5.4 kb-long ssDNA molecules in presence of the indicated proteins. (B) Proportion of ssDNA molecules with Rad51 filaments (left) and number of Rad51 segments among these positive ssDNA molecules (right). The number of segments were compared using a two-tailed Mann-Whitney test. * p-value < 0.0001. (C) Distribution of total (left) individual (center) and maximum (right) Rad51 segment length per ssDNA molecule. Conditions are color-coded as in panels A) and B). Distributions were compared using a non-parametric Kolmogorov-Smirnov test. * p-value < 0.0001.

### Stochastic modeling of disruption-driven Rad51-ssDNA filament growth

How could the Rad51 stripping activity of Srs2 be driving filament growth? We envisioned that short Rad51-ssDNA segments may act as roadblocks for each other’s growth, leading to highly segmented filaments. By turning over a subset of these segments, Srs2 may enable the growth of the surviving ones, resulting in overall longer individual segments. To gain insights into this maturation process, we developed a stochastic model of filament assembly/disassembly. We modeled a 5 kb-long ssDNA strand as a unidimensional lattice. At every time step, Rad51 segments may nucleate, grow or dissociate (see **Methods**). In particular, the filament dynamics is controlled by two main parameters (**Fig. 7A**):

**Figure 7:**
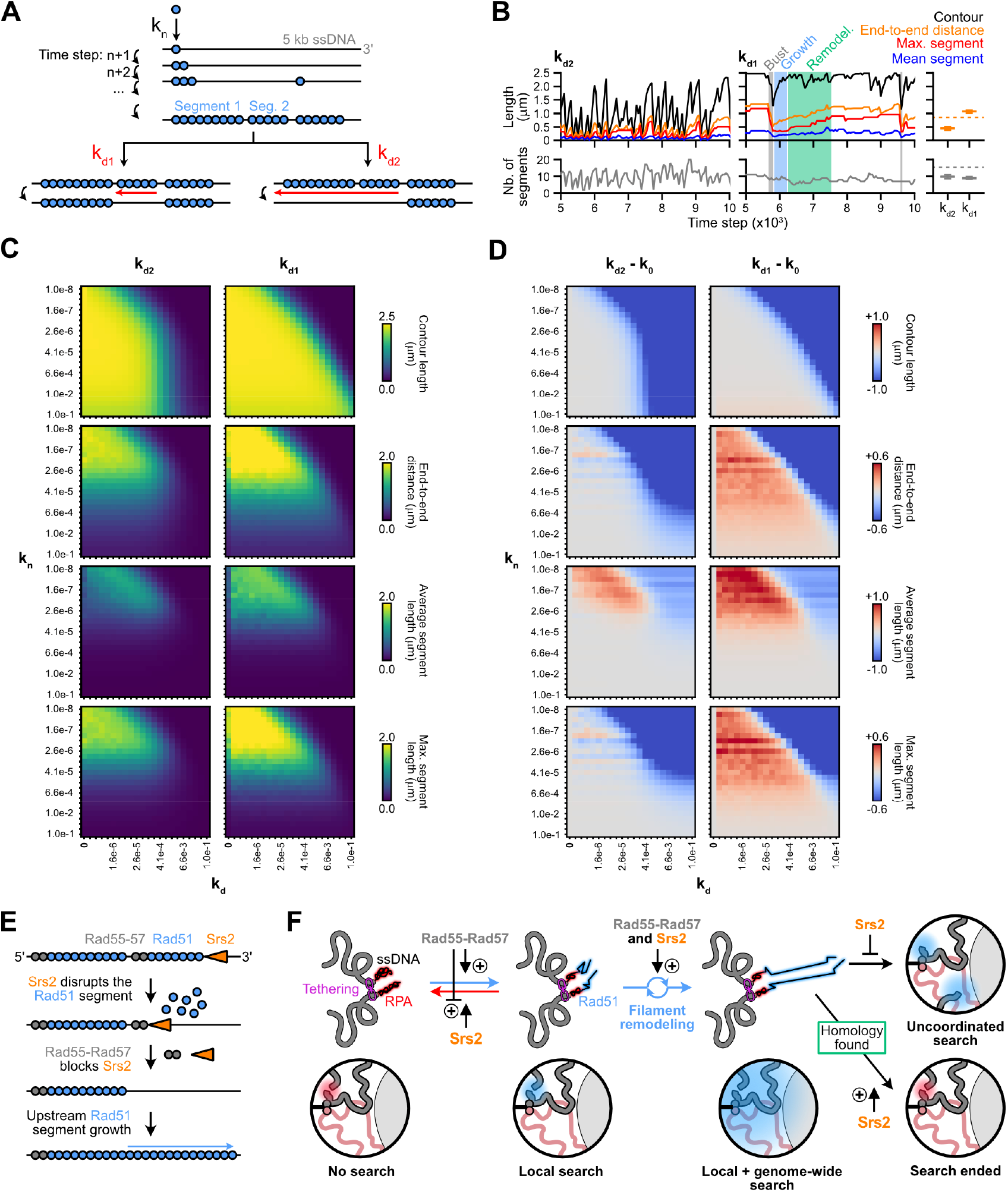
Stochastic modelling suggests a mechanism for Srs2-mediated Rad51-ssDNA filament stiffening. (A) Rationale of the Monte-Carlo simulations illustrating the two modes of Rad51 filament disassembly (k_d1_ and k_d2_). (B) Example of individual time series of Rad51 filament dynamics with parameters k_n_ =8.2×10^-5^ and k_d2_ and k_d1_ = 8.2×10^-4^. k_d1_ uniquely features remodeling phases during which long segments can grow at near maximal contour length. The aggregated steady-state values of 40 replicates are shown on the right. Dotted line: k_d_ = 0. (C) Heatmaps showing the steady-state Rad51 filament contour length, end-to-end distance, average segment length, and maximum segment length per ssDNA molecule with varying k_n_, k_d1_ and k_d2_. (D) Effect of k_d1_ and k_d2_ (relative to no k_d_) on the steady-state Rad51 filament contour length, end-to-end distance, average segment length, and maximum segment length per ssDNA molecule. From data in C). (E) Model of the interplay between Rad55-Rad57 and Srs2 in promoting Rad51-ssDNA filament remodeling. (F) Model for the individual and joint roles of Rad55-Rad57 and Srs2 in regulating homology search onset, reach, coordination and inactivation.

- The probability k_n,_ per position along the ssDNA to nucleate a Rad51 segment at a given time step; *i*.*e*. to add a monomer of Rad51 on DNA at a non-contiguous position relative to pre-existing Rad51 segments.
- The probability k_d_ per Rad51 segment(s) to dissociate at a given time step. This parameter controls either the dissociation of a single Rad51 segment (k_d1_), or of all the Rad51 segments 5’ from a randomly selected segment (k_d2_) (**Fig. 7A**). The k_d1_ thus mimics a situation in which Rad55-Rad57 blocks Srs2 once it reached the 5’ end of a Rad51 segment, while k_d2_ represents a situation in which Srs2 processively translocates and acts along the entire ssDNA from its loading site (Kaniecki *et al*, 2017; De Tullio *et al*, 2017; Roy *et al*, 2021).

Rad51 segments grow in the 3’ direction by a single monomer at every time step but contiguous segments cannot merge (**Fig. 7A** and **S10A**). The nucleation probability is thus relative to the filament growth rate. The steady-state filament segmentation, segment length, contour length (*i*.*e*. total ssDNA length covered by Rad51) and corresponding average end-to-end spatial distances were computed for each set of parameters determined over 8 orders of magnitude (**Fig. 7B, C** and **S10B, C**; **Methods**).

In the absence of dissociation (k_d_=0), the greatest end-to-end distances were achieved at the lowest nucleation probabilities: one or a few segments grow to cover the entire ssDNA (k_n_=10^-8^, **Fig. 7C**). Increasing k_d2_ caused either no change or a decrease in contour length and end-to-end distance (**Fig. 7C, D**). Differently, increasing k_d1_ resulted in an increase in end-to-end distance over a large parameter space (**Fig. 7C, D**). Such an increase occurred at constant contour length (**Fig. 7C, D**) and, for k_n_≤4.1×10^-5^, with minimal change to the number of segments (**Fig. S10B, C**). Most of this increase could be accounted for by the length of the longest Rad51 segment (**Fig. 7B-D**). Inspection of individual trajectories revealed remodeling phases, during which a progressive increase of the end-to-end distance was driven by dissociation of small segments, enabling the growth of the surviving upstream segment (**Fig. 7B**). The difference between k_d1_ in k_d2_ is particularly striking at high dissociation probabilities (≥1×10^-4^), in which k_d2_ clears most Rad51 segments while k_d1_ keeps promoting end-to-end distance gains (**Fig. 7C** and **S10D**). These results indicate (i) that disruption of Rad51 from ssDNA on a per-segment basis can drive the growth of surviving Rad51 segments, and (ii) that this proofreading of Rad51 filament structure promotes the formation of one or a few critically long segments sufficient to drive a substantial, albeit intermittent, increase in end-to-end distance, *i*.*e*. ssDNA stiffening (**Fig. S10E**). This simple change to Rad51 filament dissociation is thus sufficient to qualitatively recapitulate the *in vitro* context in which both Rad55-Rad57 and Srs2 are included in the Rad51 filament formation reaction, which uniquely produces long continuous Rad51 segments (**Fig. 6**). It suggests that Rad55-Rad57 promotes overall ssDNA stiffening by restricting the disruption activity of Srs2 to single Rad51-ssDNA segments (**Fig. 7E**).

## Discussion

Here we tracked the core HR steps in cells defective for the Rad51 paralogs Rad55-Rad57 and the Srs2 helicase following site-specific DSB induction: resection, end-tethering, Rad51-ssDNA and -dsDNA binding, homology search by both DSB ends, absolute levels of D-loop DNA joint molecules at two competing donors, as well as the extension of these D-loops. It both confirmed previous proposed functions and revealed new roles for these proteins in regulating homology search (i) onset, (ii) coordination between the two DSB ends, (iii) reach across the genome, and (iv) inactivation upon donor identification (**Fig. 7F**).

### Rad55-Rad57 stimulates Rad51-ssDNA filament formation and antagonizes its disruption by Srs2

The main proposed functions for Rad55-Rad57 in stimulating the formation of Rad51-ssDNA filaments and in antagonizing the Rad51-ssDNA filament disruption activity of Srs2 have been inferred from biochemical reconstitution experiments *in vitro*, the cytological analysis of foci formed by non-functional fluorescently-tagged Rad51, and by non-calibrated semi-quantitative ChIP assays (Sung, 1997; Gasior *et al*, 2001; Sugawara *et al*, 2003; Lisby *et al*, 2004; Fung *et al*, 2009; Liu *et al*, 2011; Xu *et al*, 2013; Elango *et al*, 2017; Roy *et al*, 2021; Maloisel *et al*, 2023; Deveryshetty *et al*, 2025). Here, we corroborated both of these functions using more direct methodologies to track Rad51-ssDNA filament formation by calibrated strand-specific ChIP-seq against a functional V5-tagged Rad51 protein (**Fig. 2**) as well as Rad51-ssDNA filament activity in cells by tracking homology search with ssHi-C (**Figs. 4** and **5**) and homology identification with DLC (**Fig. 1**). The delayed assembly of Rad51-ssDNA filaments in the *rad57*Δ *srs2*Δ mutant relative to WT or *srs2*Δ cells indicates that Rad55-Rad57 indeed stimulates their formation, consistent with the conserved chaperone role proposed for Rad51 paralogs complexes (Roy *et al*, 2021; Belan *et al*, 2021; Deveryshetty *et al*, 2025; Greenhough *et al*, 2023, 2026; Koo *et al*, 2026; Rawal *et al*, 2026). The more severe defects in Rad51-ssDNA filament assembly and function in the *rad57*Δ than in the *rad57*Δ *srs2*Δ mutant shows that Rad55-Rad57 antagonizes the Rad51-ssDNA filament disruption activity of Srs2. It cannot be straightforwardly deduced, from these *in vivo* data, whether Rad55-Rad57 physically antagonizes Srs2 in addition to promote reassembly of the Rad51-ssDNA filament (Liu *et al*, 2011; Roy *et al*, 2021). However, the Rad51-dependent stiffening of ssDNA in cells and the structural changes to Rad51-ssDNA filaments imparted by Rad55-Rad57 and Srs2 *in vitro* suggest that Rad55-Rad57 can directly block Rad51-ssDNA filament disruption by Srs2, in addition to promote filament assembly (see below).

### Rad55-Rad57 and Srs2 together promote distant homology search by remodeling Rad51-ssDNA filament structure

Mapping of Rad51-ssDNA contacts genome-wide (**Figs. 4** and **5**) and quantification of D-loops formed at competing intra- and inter-chromosomal donors (**Fig. 1B, C**) reveal that the *rad57*Δ *srs2*Δ mutant is specifically defective in homology search at distant chromosomal regions, despite a recruitment of Rad51 at the DSB close to that observed in WT cells (**Fig. 2**). Rad55-Rad57 and Srs2 together thus promote genome-wide homology search, a process that depends on the assembly of long, rigid Rad51-ssDNA filaments in *S. cerevisiae* (**Fig. 7F**) (Liu *et al*, 2023; Dumont *et al*, 2024; Piazza & Taddei, 2026). Accordingly, purified Rad55-Rad57 and Srs2 reduced the segmentation of Rad51-ssDNA filaments, resulting in the formation of long continuous Rad51 segments *in vitro* (**Fig. 6**). This remodeling increased the overall stiffness of ssDNA, providing the molecular underpinnings for the ssDNA stiffening inferred from the distribution of ssDNA contacts (**Fig. 5D-F**). Stochastic modeling indicated that the disruption activity of Srs2 could lead to such an outcome if it was restricted to individual Rad51-ssDNA segments, relieving an impediment for the growth of upstream segments (**Fig. 7E** and **S10E**). We propose that Rad55-Rad57 achieves such a restriction by capping the 5’ end of the individual Rad51 segments it contributed to nucleate (**Fig. 7E**). This model additionally explains how Rad51 paralogs complexes at the 5’ end of Rad51 segments can overall protect segmented Rad51-ssDNA filaments from disruption by Srs2, a helicase with 3’-to-5’ directionality (Rong & Klein, 1993). The 5’ filament association of the human RAD51 paralogs complexes (Greenhough *et al*, 2026; Koo *et al*, 2026; Rawal *et al*, 2026) and the demonstrated or predicted 3’-to-5’ directionality of the helicases endowed with hRAD51 stripping activity (*i*.*e*. RECQL5, FBH1, and PARI) (Hu *et al*, 2007; Moldovan *et al*, 2012; Simandlova *et al*, 2013) suggests a broad conservation of this Rad51-ssDNA filament remodeling mechanism.

The whole length of the ssDNA need not to belong to a continuous Rad51-ssDNA segment for DSB-proximal sites to be projected into a distant nuclear area. Indeed, by virtue of its high stiffness, a single 1 kb-long Rad51 segment will stretch ssDNA over ∼500 nm, driving the spatial separation of DSB-distal and DSB-proximal ssDNA sites (**Fig. 7F**). Hence, the stochastic local remodeling of initially segmented Rad51-ssDNA filaments only needs to result in the growth of a single Rad51 segment anywhere along the resected ssDNA to enable distant homology search (**Fig. 7E**). This remodeling strategy allows for a fast onset of local homology search by initially highly-segmented Rad51 filaments while also providing the means to subsequently transition to a genome-wide search (**Fig. 7F**). The per-segment dissociation rate tunes both the rate of this transition and the duration of a distant search event. Such a model, in which only a subset of Rad51 molecules are part of the longest segment, may account for the abrupt contraction events observed cytologically without major changes to the overall fluorescence intensity of the Rad51 structure (Liu *et al*, 2023).

### Srs2 promotes coordinated search by Rad51-ssDNA filaments on both DSB ends

DSB end-tethering promotes coordinated search between filaments on both sides of the DSB (Dumont *et al*, 2024). Functionally, this coordination likely stimulates HR repair, as tethering of non-cognate ends causes pronounced kinetics delay in HR product formation (Jain *et al*, 2016). The maintenance of DSB end-tethering over the course of HR results from multiple handovers: the initial tethering by Sae2-MRX is subsequently passed on to Exo1 and the Ddc1-Mec3-Rad17 (human 9-1-1) clamp (Kaye *et al*, 2004; Lobachev *et al*, 2004; Clerici *et al*, 2005; Nakai *et al*, 2011; Piazza *et al*, 2021; Phipps *et al*, 2024). These determinants indicate that a main tethering point between the two DSB ends during HR repair is located at the resection front. Donor identification, in our system where homology exists to only one DSB end, also led to a modest loss of DSB end-tethering, which could be aggravated in the absence of PCNA/Pol*δ* (**Fig. S6D-E**) (Dumont *et al*, 2024). These observations indicated that end-tethering was sensitive to the HR status downstream of homology search, requiring concomitant homology identification by both ends. Here, we further revealed that DSB end-tethering could be compromised in cells deficient for the ssDNA translocase activity of Srs2, which undermined homology search coordination between DSB ends (**Fig. 4G**). This progressive loss of end-tethering depended on Rad51 and Rad57 but not on the presence of a homologous donor or of PCNA/Pol*δ*, which suggests that Rad51-ssDNA filaments formed in the absence of Srs2 and/or a dysregulated pre-synaptic activity of these filaments, undermine DSB end-tethering.

Direct and indirect methods to visualize DSB-induced Rad51 filaments revealed single filament structures that can occasionally branch in yeast and mammalian cells (Liu *et al*, 2023; Sharma *et al*, 2024; Friskes *et al*, 2025; Taniguchi *et al*, 2025). Despite the limitation in spatial resolution of conventional fluorescence microscopy, these observations are suggestive of a predominantly longitudinal association of Rad51 filaments (Piazza & Taddei, 2026), raising the possibility of contact points along Rad51-ssDNA filaments. Indeed, recent *in vitro* studies revealed the longitudinal bundling of human RAD51 filaments mediated by the TR2 domain of BRCA2 and by RAD52 (Appleby *et al*, 2023; Alshareedah *et al*, 2026). Paired hRAD51 filaments were also reported to form spontaneously in the presence of ADP *in vitro* (Luo *et al*, 2023). Two types of interactions may thus contribute to DSB end-tethering during homology search: those involving factors present at the resection edges, and others along Rad51-ssDNA filaments. The precise molecular nature of these links in cells, and of the mechanism driving the loss of end-tethering in contexts in which the metabolism of Rad51-ssDNA filaments is perturbed in Srs2-deficient cells, remain to be established.

### Srs2 inactivates homology search following homology identification

Whether homology identification feedbacks onto the search process, conducted in parallel along Rad51-ssDNA filaments, or whether homology search termination merely results from the elimination of the recombinogenic substrate upon repair completion, has not been addressed. Here, by quantifying D-loops in a system that does not authorize formation of a dsDNA repair product (**Fig. 1**), we provided evidence that the competence of ssDNA to form D-loops is compromised post-homology identification. This reduction coincides with D-loop extension, requires PCNA/Pol*δ* (Reitz *et al*, 2023) and, as we show here, Srs2 (**Fig. 1B**). Consequently, inactivation of homology search occurs subsequently to D-loop extension, presumably upon SUMO-PCNA-mediated recruitment of Srs2 (Papouli *et al*, 2005; Pfander *et al*, 2005; Liu *et al*, 2017; Arbel *et al*, 2020). From that point, translocation of Srs2 in the 3’-to-5’ orientation is expected to dissociate Rad51 from upstream ssDNA, *de facto* terminating homology search by that ssDNA (**Fig. 2D**). Accordingly, Srs2 eliminated Rad51 from the neo-synthesized DNA generated upon D-loop extension (**Fig. 2C**). These observations thus identify an unanticipated control point of HR mediated by Srs2, whereby homology search is inhibited in the time frame separating D-loop extension from repair completion. This regulation may contribute to Srs2’s role in suppressing the formation of toxic HR intermediates (Gangloff *et al*, 2000) as well as multi-invasion DNA joint molecules and repeat-mediated chromosomal rearrangements (Putnam *et al*, 2009, 2016; Elango *et al*, 2017; Piazza *et al*, 2017).

### Srs2 inhibits the first strand synthesis of BIR

BIR is a conservative DNA synthesis pathway occurring in two steps: a first displacement DNA synthesis from the invading strand occurring in the context of a D-loop, and a second strand synthesis primed once a telomere has been captured, using the neo-synthesized DNA has a template (Donnianni & Symington, 2013; Saini *et al*, 2013; Thakre *et al*, 2026). Despite its extended D-loop disruption activity, Srs2 has surprisingly been described as a pro-BIR factor (Ruiz *et al*, 2009; Elango *et al*, 2017; Uribe-Calvillo *et al*, 2022); a proposal that originates from molecular assays that detect the final dsDNA BIR product or from genetic assays that require cell survival following BIR completion. Here, by tracking Rad51 binding to ssDNA, we show that Srs2 inhibits the rate of the first strand synthesis of BIR by about ∼20 kb/hr (**Fig. 2C**). This observation is consistent with Srs2’s ability to disrupt extended D-loops *in vitro*, to cause a reduction in D-loop levels in *S. cerevisiae* cells (Liu *et al*, 2017; Piazza *et al*, 2019; Hung *et al*, 2025), and of its *S. pombe* homologs to promote template-switch at stalled replication forks (Jalan *et al*, 2019). We thus propose that the pro-BIR role of Srs2 arises from additional functions downstream of first strand synthesis, such as inactivating the DNA damage checkpoint elicited by the presence of extensive RPA-coated ssDNA (Vaze *et al*, 2002; Dhingra *et al*, 2021), and/or by promoting second strand synthesis (Saini *et al*, 2013; Donnianni & Symington, 2013). These late effects may have masked the upstream role of Srs2 in inhibiting the first strand synthesis when scoring BIR product alone, highlighting the value of tracking individual HR intermediates.

### Limitations of the study

First, our inability to disentangle the contribution of ssDNA stiffening by Rad51 from that of a loss of end-tethering in promoting distant homology search complicated the interpretation of the defects observed at the D-loop level in the *srs2*Δ mutant. Second, the DSB-distal and DSB-proximal ssDNA contact distributions obtained in the *srs2*Δ mutant could not be satisfyingly fitted with a single *L*_*p*_ value (**Fig. 5D, E**), which suggested that ssDNA stiffening was not uniform in this mutant. Our population-based approaches (ssChIP-seq and ssHi-C) cannot resolve this heterogeneity. These limitations prevented us from establishing the contribution of Srs2 alone in Rad51-ssDNA filament stiffening and competence for genome-wide homology search. The intra/inter donor preference determined with DLC at 2 hours (**Fig. 1C**), when end-tethering defects and loss of search coordination are minimal (**Fig. 3A** and **4F**), suggested a specific defect in genome-wide homology search in Srs2-deficient cells. Finally, the sensitivity of D-loop Capture, which can detect ∼1 D-loop in 10^4^ cells (Reitz *et al*, 2022; Djeghmoum & Piazza, 2025), remains insufficiently sensitive to confidently quantify D-loops formed at the inter donor in the *rad57*Δ mutant.

## Methods

### Experimental procedures

#### Saccharomyces cerevisiae strains

The genotype of *Saccharomyces cerevisiae* (W303

*RAD5*+ background) strains used in this study are listed in **Table S1** and the annotated sequences of the relevant genetic constructs are available in **Dataset S1**. The mutagenesis of the endogenous HO cut-site at *MAT* (chr. III), the DSB-inducible construct at *ura3* (chr. V), the inter-chromosomal donor at *lys2* (chr. II) and the intra-chromosomal donor at *can1* (chr. V) have been described previously (Piazza *et al*, 2018, 2019, 2021; Djeghmoum & Piazza, 2025). Briefly, the DSB-inducible construct contains a 327 bp-long fragment of the PhiX phage genome flanked by multiple restriction sites (including *Eco*RI and *Hind*III) making a unique 453 bp sequence pad used in the DLC assay (below), a 2,086 bp-long sequence corresponding to the first half of the *LYS2* gene (+4 to +2,090), and a 117 bp HO cut-site. It is positioned on chr. V at position 116,167 in S288c coordinates, replacing the *URA3* gene. The intra-chromosomal donor is located on chr. V at position 33,027 (*can1*), with an *Eco*RI site used for D-loop Capture present 233 bp upstream, and a *HindIII* site used for quantifying D-loop extension present 138 bp downstream. The inter-chromosomal donor is located on chr. II at position 468,743 (*lys2*) on the Crick strand, with an *Eco*RI site used for D-loop Capture present 165 bp upstream, and a *HindIII* site used for quantifying D-loop extension present 375 bp downstream.

The introduction of the V5 tag between Gly54 and Gly55 of Rad51 at its endogenous locus has been performed by CRISPR/Cas9 gene-targeting with the guide 5’-tggcggattgcaggagcaag-3’ and a dsDNA repair template annotated in **Dataset S1**. The V5 tag is flanked by 16 amino acids-long flexible linkers on each side. The annotated sequence of the resulting *iV5-RAD51* genetic construct is available in **Dataset S1**. The *trp1::pGAL1-HO* construct for over-expressing the HO endonuclease has been described previously (Pannunzio *et al*, 2008). The *his3::pADH1-OsTIR1-9Myc::HIS3* construct for expressing the OsTir1 E3-Ubiquitin ligase and the Scc1-(Pk3-)AID constructs have been described previously (Dauban *et al*, 2020; Piazza *et al*, 2021). The *rad51::kanMX, srs2::KanMX, rad55::natMX*, and *rad57::LEU2* gene deletions were obtained upon short homology ends-out gene targeting of PCR products amplified from Longtine vectors (Longtine *et al*, 1998). The *srs2-K41A* point mutant from ref. (Piazza *et al*, 2019) has been generated by pop-in/pop-out. The Rfc1-iAID and Pol3-iAID constructs have been obtained by crossing against strains generated in ref. (Donnianni *et al*, 2019). Genetic alterations were verified by PCR and Sanger sequencing. The correct genotype was further verified from bam alignment files of ChIP-seq, Hi-C and/or ssHi-C experiments.

#### Culture media

Liquid YPD (1% yeast extract, 2% peptone, 2% dextrose) and YEP-lactate (1% yeast extract, 2% peptone, 2% lactate) media were prepared according to standard protocols (Treco & Lundblad, 2001) using ultrapure milli-Q water. YPD plates containing campthotecin (Euromedex cat. TO-C146) and methyl methane sulfonate (Sigma-Aldrich cat. 129925) were prepared within a day prior to usage.

#### Induction of DSB formation

Site-specific DSB induction upon HO over-expression in an asynchronous cell population was performed as described previously (Piazza *et al*, 2018, 2019). Briefly, an isolated colony was grown overnight with agitation in YPD at 30°C and diluted in YEP-lactate media. HO expression was induced by addition of 2% galactose to exponentially growing cultures (∼5.10^6^ cells/mL).

Site-specific DSB induction upon HO over-expression in cells synchronously released in S-phase was performed as described previously (Dumont *et al*, 2024). Briefly, exponentially growing cells in YEP-lactate (30°C) were synchronized at the G1/S transition by addition of 1 µg/ml alpha-factor (GeneCust) every 30 minutes for 3-4 hours. Cells were washed 3 times with 50 mL of pre-warmed YEP-lactate and released in S-phase in YEP-lactate supplemented with 2% galactose. The quality of the synchrony was verified by flow cytometry.

Scc1-(Pk3-)AID depletion was induced at the time of DSB induction in cells synchronously released from a G1 block upon addition of 2 mM 3-Indolacetic Acid (IAA, aka auxin), as previously described (Dumont *et al*, 2024). Rfc1-iAID and Pol3-iAID were co-depleted upon addition of 50 μg/mL doxycycline to repress their transcription and 1.5 mM IAA 1 hour after S-phase release and DSB induction, as previously described (Dumont *et al*, 2024).

#### Flow cytometry

Approximately 10^7^ cells were collected by centrifugation, re-suspended in 70% ethanol and fixed at 4°C for at least 24 hours. Cells were pelleted, washed in 1 mL of 50 mM Sodium citrate pH 7.0, and resuspended in 1 mL 50 mM Sodium citrate pH 7.0. 200 µL of the resuspension was treated with 200 μg RNase A (Euromedex, cat. 9707-C). Following incubation at 50°C for 1 hour or 37°C overnight, cells were pelleted and resuspended in 1.8 mL of 50 mM Sodium citrate pH 7.0 with 2 µM of SYTOX™ Green (Thermo Scientific, S7020), incubated for 1 hour at room temperature in the dark, and sonicated for 10 seconds on a Bioruptor. Flow cytometry profiles were obtained on a MACSQuant machine and analyzed using Flowing Software 2.5.1.

#### D-loop Capture assay

D-loop capture assay was performed as described in (Reitz *et al*, 2022; Djeghmoum & Piazza, 2025; Piveteau *et al*, 2026), except for data in **Fig. S2** performed as in (Piazza *et al*, 2019), which omitted psoralen crosslink reversal prior to chimera quantification by qPCR. Briefly, a site-specific DSB was induced upon over-expression of the HO endonuclease by adding galactose at a final concentration of 2% to an exponentially growing cell culture in YEP-lactate media. 2.10^8^ cells we collected prior to, and at various time points after galactose addition. For DLC, cells were re-suspended in crosslinking solution (0.1 mg/mL Trioxsalen (Sigma-Aldrich T6137), 50 mM tris HCl pH 8.0, 50 mM EDTA, 20% ethanol) and the DNA was crosslinked with ∼32 J/cm^2^ UV-A (365 nm) irradiation in a Bio-link – BLX365 (Vilber-Lourmat, cat. 611110831) with permanent orbital agitation (∼50 rpm). Cell were spheroplasted for 15 min at 37°C with 3.5 μg/mL Zymolyase 100T in spheroplasting buffer (0.4 M sorbitol, 0.4 M KCl, 40 mM phosphate buffer pH 7.4, 0.5 mM MgCl_2_) and washed twice with spheroplasting buffer and three times with 1X Cutsmart buffer (20 mM tris acetate pH 7.9, 50 mM potassium acetate, 10 mM magnesium acetate, 100 μg/mL BSA) at 4°C. Pellets were resuspended in 1.4X Cutsmart buffer, flash frozen in liquid nitrogen and stored at -70°C. Cells were lysed upon addition of 0.1% SDS at 65°C for 10-15 min in the presence of a 80mer oligonucleotide (APO563; **Table S2**), whose annealing restores the EcoRI restriction site on the resected broken molecule. DNA was recovered from the spheroplasts, digested by EcoRI-HF (NEB, cat. R3101L), and ligated with T4 ligase (NEB, cat. M0202) at low concentration (∼1.8 × 10^4^ genome/µl). DNA was extracted with phenol-chloroform after protein degradation using proteinase K. Psoralen inter-strand crosslinks and adducts was reversed in 100 mM KOH at 90°C for 30°C. The pH was neutralized upon addition of 66 mM of NaoAc pH 5.2. Approximately 6 × 10^5^ genome equivalent were used per quantitative PCR (qPCR) reaction, performed in duplicate, on a CFX96 Touch Deep Well Real-Time PCR Detection System (Bio-Rad cat. 3600037), using the SsoAdvanced Universal SYBR Green Supermix (Bio-Rad, cat. 1725274), following manufacturer’s instructions. Primers used are listed in **Table S2**. qPCR analysis were performed as described in (Reitz *et al*, 2022) using Bio-Rad CFX Maestro and Microsoft Excel.

#### Calibrated strand-specific chromatin immunoprecipitation and sequencing (ChIP-Seq)

Calibrated strand-specific ChIP-Seq was performed by combining protocols from (Hu *et al*, 2015) and (Peritore *et al*, 2021). Approximately 15 OD_600_ of a culture of *Saccharomyces cerevisiae* (∼1.5×10^8^ cells) were mixed with 0.75 OD_600_ of an exponentially growing culture of *Candida glabrata* cells (KN23308) expressing Scc1-V5, in triplicate. Each triplicate reaction was crosslinked in a final volume of 37 mL with 3% formaldehyde (Sigma-Aldrich cat. F8775) for 30 minutes with orbital agitation at 120 rpm. Formaldehyde was quenched with 330 mM Glycine for 20 minutes with orbital agitation at 120 rpm, washed twice at 4°C with 1X PBS, and the pellets snap frozen in liquid nitrogen and stored at - 70°C.

Mechanical cell lysis was performed using Precellys in 2 mL tubes (3 rounds at 6,800 rpm during 12 seconds - 45 seconds in ice) in 300 μL of lysis solution (50 mM HEPES-KOH pH8, 140 mM NaCl, 1 Mm EDTA, 1% Triton X100, 0.1% Sodium deoxycholate, 1 mM PMSF and a protease inhibitor cocktail (Sigma-Aldrich, cat. 11836170001). The lysate was collected and lysis solution was added to reach 1 mL. The lysate was sonicated on a Covaris S220 to shear DNA into 200-1,000 bp fragments in 1 mL tube (Covaris, cat. 520130). The lysate was clarified twice by centrifugation at 10,000 g at 4°C. 27 μL were collected per triplicate, pooled to make the input, and mixed with 40 μL of TES3 buffer (50 mM Tris HCl pH8, 10 mM EDTA, 3% SDS). The remainder of the clarified lysate was incubated with 2 µg of Mouse anti-V5 monoclonal antibody (Invitrogen, cat. R96025) overnight at 4°C on a rotating wheel. Immunoprecipitation was performed upon addition of magnetic Protein G Dynabeads (Invitrogen, cat. 10003D) and incubation for 2 hours at 4°C on a rotating wheel. Beads were washed with lysis solution, lysis solution with 500 mM NaCl, washing solution (10 mM Tris HCl pH8, 500 mM LiCl, 1 mM EDTA, 0.5% Igepal CA-630, 0.1% Sodium deoxycholate, 1 mM PMSF),TENa buffer (10 mM Tris-HCl pH8, 1 mM EDTA, 50 mM NaCl), resuspended in 120 μL of TES1 buffer (50 mM Tris-HCl pH8, 10 mM EDTA, 1% SDS), and incubated 15 minutes at 65°C and recover supernatant to obtain the immunoprecipitated sample (IP). The triplicate IPs and the input were decrosslinked overnight at 65°C. Following RNA and protein digestion, DNA was purified by 25:24:1 Phenol:Chloroform:Isoamyl alcohol (Sigma-Aldrich, cat. P3803) extraction, precipitation, and resuspended in 30 μL 10 mM Tris HCl pH8 0.5 mM EDTA. The DNA concentration was quantified quantified using the Qubit DNA high sensitivity kit (Thermo Scientific, cat. Q32851) on a Qubit 2 fluorometer (Thermo Scientific, cat. Q32866). The IP and input sequencing libraries were prepared using the xGen ssDNA & low-input DNA Library Prep kit (IDT, cat. 10009859) following manufacturer’s instructions. The final library was PCR amplified for 12 and 9 cycles for the IP and input, respectively. The DNA concentration quantified on a Qubit 2 fluorometer and the library quality control, paired-end sequencing (2×150 bp) on Illumina NovaSeq6000 or NovaSeq X Plus, and data QC were performed by Novogene UK.

#### Hi-C and ssHi-C procedures

Hi-C and ssHi-C were conducted as described in (Dumont *et al*, 2024). Briefly, ∼1.5×10^9^ haploid cells were fixed with 3% formaldehyde (Sigma-Aldrich, cat. F8775) for 30 minutes at RT with orbital agitation at 120 rpm. Formaldehyde was quenched with 330 mM glycine for 20 minutes at RT with orbital agitation at 120 rpm. Cells were washed twice with cold water at 3,000 g for 10 minutes. Pellets were split in two tubes of ∼100 mg and frozen at -80° C.

Cell pellets were thawed in ice, resuspended in 10 mL of Zymolyase solution (0.25 mg/mL Zymolyaze 100T (Carl Roth, cat. 9329), 1 M D-sorbitol (Sigma-Aldrich, cat. S6021), 50 mM Tris-HCl pH 7.5, 10 mM 2–mercaptoethanol (Sigma-Aldrich, cat. M3148)) and spheroplasted at 210 rpm for 18 minutes at 30° C (cells spheroplasting is checked under the microscope and the incubation time adjusted accordingly). Spheroplasts are pelleted at 3,000 g for 10 minutes at 4° C, washed with 4 mL cold 1X PBS, and resuspended in a fresh EGS crosslinking solution (3 mM ethylene glycol bis(succinimidyl succinate) (Fisher Scientific, cat. 10350924) in 1X PBS), and incubated at 30° C for 40 minutes at 100 rpm. EGS was quenched with 400 mM glycine at RT for 5 minutes at 100 rpm. The crosslinked cellular material was pelleted at 3,000 g for 10 minutes at 4° C and washed twice in cold 1X PBS supplemented with anti-protease (Sigma-Aldrich, cat. 11836170001). Pellets were snap-frozen in liquid nitrogen and stored at - 80°C.

The crosslinked pellets were thawed in ice and resuspended in cold H_2_O supplemented with anti-protease. Between 2 and 5.10^7^ cells were used for the remainder of the procedure. The ssHi-C procedure uses the Arima Hi-C kit (Arima Genomics, cat. A410079), which employs a dual restriction digestion (DpnII and HinfI) that yields a median fragment length of 108 bp in *S. cerevisiae*. The protocol follows manufacturer’s instructions with the following modification: after the lysis step, the pool of 80 nt-long, 3’-blocked, PAGE-purified annealing oligonucleotides (stock concentration 100 nM each; **Table S2**) is added at a final concentration of 7 nM each. Annealing occurs during the subsequent incubation at 62° C for 10 minutes.

DNA was fragmented into 300-400 bp fragments using Covaris M220 sonicator. Preparation of the libraries for paired-end sequencing on an Illumina platform was performed using the Thermofisher Collibri ES DNA Library Prep Kit for Illumina Systems with UD indexes (cat. A38606024) following manufacturer’s instructions. The library was amplified in triplicate PCR reactions using oligonucleotides corresponding to the Illumina sequence adaptors (5’-AATGATACGGCGACCACCGAGATCTACAC-3’ and 5’-CAAGCAGAAGACGGCATACGAGAT-3’) and Phusion DNA polymerase (New England Biolabs, cat. M0531) for 11 cycles. PCR products were purified with AMPure XP beads (Beckman-Coulter, cat. A63881) and resuspended in pure H_2_O. The Hi-C library is quantified using the Qubit DNA high sensitivity kit (Thermo Scientific, cat. Q32851) on a Qubit 2 fluorometer (Thermo Scientific, cat. Q32866).

The ssHi-C library consists in an enrichment of ssDNA contacts from the Hi-C library using a panel of biotinylated “capture” oligonucleotides targeting the fragments targeted by annealing oligonucleotides as well as 7 control dsDNA sites (**Table S2**) (Dumont *et al*, 2024). 500 ng of PCR products were saturated with 2 μg of unlabeled Cot1 human DNA (Invitrogen, cat. 15279-011) and 0.2 nmol of oligonucleotides corresponding to the universal Illumina adaptors (see above) and dehydrated at 60° C for 1 hour in a Speedvac (Thermo Scientific, cat. DNA130-230). DNA was resuspended in 1X xGen Hybridization buffer and 0.15X of xGen Hybridization Enhancer (Integrated DNA Technologies, cat. 10010352), and the pool of biotinylated capture oligonucleotides (stock 1.8 μm each) at a final concentration of 0.6 μM. DNA was denatured at 95° C for 5 minutes, and annealing of capture oligonucleotides to their target occurred at 65° C for 4 hours. Biotin pulldown on streptavidin-coated dynabeads C1 (Invitrogen, cat. 65001) was performed according to manufacturer’s instructions, and DNA was recovered in pure H_2_O. PCR amplification (10 cycles) and DNA purification on AMPure XP beads was carried out as before. The capture procedure was repeated once and the final library was amplified for only 8 cycles. PCR products (300 ng) were digested at 37° C for 1 hour with 10 units of MfeI-HF and SspI-HF in 1X rCutsmart buffer (New England Biolabs, cat. R3589, R3132 and B7204, respectively) and the enzymes heat-inactivated at 65° C for 20 minutes. This digestion step ensures elimination of the chimeric molecules involving the fragments of interest (*i*.*e*. targeted by annealing and capture oligonucleotides) that were in a dsDNA form in the cell population at the time of crosslinking, leaving only ssDNA contacts at these sites (**Fig. 4A**). DNA was purified with AMPure XP beads, resuspended in pure H_2_0. The ssHi-C library was quantified using the Qubit DNA high sensitivity kit (Thermo Scientific, cat. Q32851) on a Qubit 2 fluorometer (Thermo Scientific, cat. Q32866). The Hi-C and ssHi-C library quality control, paired-end sequencing (2×150 bp) on Illumina NovaSeq6000 or NovaSeq X Plus, and data QC were performed by Novogene UK.

#### Protein extraction and Western blotting

Protein extracts for western blot were prepared from 5×10^7^ to 10^8^ cells. Cells were lysed in cold NaOH buffer (1.85 N NaOH, 7.5% v/v beta-Mercaptoethanol) for 10 min in ice. Addition of trichloroacetic acid (15% final) for 10 min in ice allowed protein precipitation. After centrifugation at 15,000 g for 5 min, the pellets were resuspended in 100 µL of SB++ buffer (180 mM Tris-HCl pH 6.8, 6.7 M Urea, 4.2% SDS, 80 µM EDTA, 1.5% v/v Beta-mercaptoethanol, 12.5 µM Bromophenol blue). Denaturation was performed by heating 5 min at 65°C. Pre-cleared extracts were resolved on 12% precast polyacrylamide gel (Bio-Rad, cat. 4561043) and blotted on a PVDF membrane (GE Healthcare, cat. 10600023). Membranes were probed with mouse anti-AID antibody diluted at 1:1000 (clone 1E4, MBL life science, M214-3) or anti-GAPDH antibody diluted at 1:10000 (clone GA1R, Invitrogen, MA5-15738), and revealed with an HRP-conjugated rabbit anti-mouse IgG antibody diluted at 1:5000 (Invitrogen, A16160) using Immobilon Forte western HRP substrate (Merck, WBLUF0100) and a Chemidoc MP Imaging system (BioRad).

#### Protein purification

Yeast proteins Rad51, RPA, Srs2, and Rad55-Rad57 were purified following established protocols (Sung, 1994; Papouli *et al*, 2005; Binz *et al*, 2006; Liu *et al*, 2011). Specifically, Rad51, RPA, and Rad55-Rad57 were isolated from yeast cells. For native Rad51, the clarified lysate underwent ammonium sulfate precipitation, followed by sequential chromatography on Q Sepharose, hydroxyapatite, and Mono Q columns to isolate the respective fractions containing Rad51. Native RPA was purified by loading the clarified lysate onto Affi-gel Blue and Mono Q columns, with elution facilitating the collection of RPA-containing fractions. GST-Rad55-His6-Rad57 was isolated via sequential affinity chromatography on Glutathione Sepharose 4B and Ni-NTA agarose columns. His6-Srs2 was purified from Sf9 cells infected with recombinant baculovirus, with the cell extract subjected to sequential chromatography on SP Sepharose, Ni-NTA agarose, and Mono Q columns. The peak fractions obtained from each purification step were concentrated, rapidly frozen in liquid nitrogen, and stored at -80°C.

#### Rad51-ssDNA filament assembly in vitro and imaging

To assemble the protein–DNA filament, 2.34 μM Rad51 in the presence or absence of 0.43 μM Rad55– Rad57 was incubated with 7 μM of a 5,400 nucleotide-long phiX174 ssDNA strand for 10 minutes at 30°C in a buffer comprising 20 mM triethanolamine (pH 7.5), 4 mM magnesium acetate, 1 mM DTT, and 3 mM ATP. Subsequently, 0.21 μM RPA was added and the mixture was incubated for an additional 10 minutes. Finally, either 0.4 μM Srs2 or a buffer control was incorporated, followed by a 10-minute incubation. The reaction mixtures were then diluted twenty-fold in a solution of 10 mM Tris-HCl (pH 7.5), 50 mM NaCl, and 5 mM MgCl2, without the use of chemical fixation. The samples were subsequently adsorbed onto 400 mesh carbon-coated copper grids (Ted Pella), negatively stained with 2% (w/v) uranyl acetate, blotted, and air-dried. Grids were imaged using a JEOL JEM-1230 transmission electron microscope. Rad51-filament images were randomly acquired from various regions on the grid. For each condition, 3 grids were utilized. Images were captured at a nominal magnification of X40,000, following minimum dose protocols, with a Tietz 2,048 × 2,048 pixel CCD camera (TVIPS, Germany).

### Data analysis procedures

#### Calibrated ChIP-seq analysis

Calibrated sequencing read mapping and analysis was performed using Tinymapper (Serizay, 2025). Briefly, both the IP and input reads were sequentially aligned using Bowtie2 on the *C. glabrata* (CBS138) and the *S. cerevisiae* (S288c R64-2-1 assembly) reference genomes to select reads that align uniquely on each genome. The ORI factor (WCE_Cg_IP_Sc_ / WCE_Sc_IP_Cg_, in which WCE_Cg_ and IP_Cg_ correspond to the number of paired reads that mapped uniquely on *C. glabrata* genome, and conversely for *S. cerevisiae* reads) was determined and ChIP-seq profiles in BigWig format were normalized per million sequences and multiplied by the ORi factor. The strand bias was computed using the bamCoverage tool of Deeptools (Ramírez *et al*, 2014). Coverage and strand bias plots were generated with custom Python scripts using the pyBigWig tool of Deeptools and Matplotlib.

#### Hi-C read alignment and data filtering

Paired-end reads in fastq.gz format were aligned and processed into a contact matrix in Graal format using the *pipeline* function of Hicstuff version 3.1.2 or 3.2.4, which uses Bowtie2 (Langmead & Salzberg, 2012) for alignment and Pairtools (Open2C *et al*, 2024) for assigning reads to restriction fragments. The pipeline was run with the following arguments: *-m* cutsite *-e* DpnII,HinfI *-q* 20 *-n -p -d -f -F -D* (*-M* graal). Briefly, paired fastq reads were pre-digested at ligated DpnII and/or HinfI sites (*-m* cutsite argument), and aligned independently using Bowtie2 in its most sensitive mode (--very-sensitive-local) to the *S. cerevisiae* reference genome R64-2-1 deleted for the *HML, MAT, HMR, URA3*, and *LYS2* sequences (S288c_DSB). Alternatively, Parasplit was used to digest the reads prior to alignment with the “--borderless” option activated. Each uniquely-mapped read with a quality equal or greater to 20 was assigned to a restriction fragment generated from the genome fasta file. Spurious contacts (*i*.*e*. uncuts and loops) were filtered as described in ref. (Cournac *et al*, 2012) using the *-f* argument and the distribution of retained events plotted using the *-p* argument. PCR duplicates were discarded with the *-D* argument. The argument *-d* was set to compute the per-chromosome probability of contact P_c_ as a function of the genomic distance *s* from pairs files. The resulting sparse matrix in Graal format was binned at 1 kb resolution using the Hicstuff *rebin* function and converted to a cooler format with the Hicstuff *convert* function. ICE normalization of the cooler file was performed using the Cooler *balance* function (Abdennur & Mirny, 2020). Iterative coarsening and balancing were achieved using the Cooler *zoomify* function, which resulted in a .mcool file. Cooler files from biological replicates were merged with cooler *merge* function. Both Graal and cooler files were used for downstream analysis.

#### ssHi-C read alignment and data filtering

The alignment of ssHi-C paired-end reads was performed as for HiC reads using Hicstuff *pipeline* function. The S288c_DSB reference genome was modified so that the annealing oligonucleotides and their target sequence on genomic DNA were placed on two artificial chromosomes, chr_artificial_dsDNA and chr_artificial_ssDNA, respectively. The sequences corresponding to the target sequences on chr. V were replaced by Ns so that each sequence is only present once in the reference genome, resulting in the S288c_DSB_LY_Capture_artificial_v12 reference. The Graal matrix generated by Hicstuff was processed using the ssHiCstuff *pipeline* function to compute genome-wide 4C-like profiles, coverage, pile-ups and statistics for individual ssDNA sites or groups thereof (Mendiboure *et al*, 2026). The same procedure was repeated with fastq files subsampled at 4M reads (seed 100) using seqtk (Li, 2026) to mitigate the influence of PCR duplicates on quantitative comparisons between ssDNA contacts across samples.

#### 4C-like ssDNA contact profiles

4C-like profiles were plotted using the ssHiCstuff *plot* function. Alternatively, 4C-like profiles binned at 1 or 10 kb were smoothed over 5 bins with the Savitzky-Golay filter using the *signal*.*savgol* function of Scipy and plotted using Graphpad Prism.

#### Computation of ssDNA contacts with distant genomic regions

The DSB region is located ∼35 kb away from *CEN5*, in the region involved in the centromeric cluster. Consequently, certain inter-chromosomal regions (*i*.*e*. other centromeres) behave as *cis* sites, which precluded the straightforward usage of inter-chromosomal contacts as a proxy for measuring interactions with distant regions. Consequently, distant regions were defined as inter-chromosomal regions excluding 160 kb on each side of the donor site on chr. II and centromeres. The rDNA was also excluded. Despite representing 60% of the genome, only ∼5% of the contacts made by the intact DSB site is made with these distant regions in metaphase-arrested cells. The proportion of contact with these regions was computed for each ssDNA site and plotted using Graphpad Prism 10.

#### Generation of contact maps from Hi-C/ssHi-C data

Ratio maps were generated from sparse matrices in Graal or Cool format using Hicstuff *view* function, SCN or ICE normalized, log-transformed and binned. Alternatively, intra-chromosomal ratio maps were generated using Serpentine (Baudry *et al*, 2020). Briefly, matrices binned at 1 kb were subsampled to contain the same number of contacts as the least covered matrix, and 30 serpentine cycles were performed with the lower and higher coverage thresholds set to 10 and 50, respectively. The ratio maps were plotted with a trend set to the mean.

#### Computation of the average contact frequency as a function of genomic distance

Computation of the contact probability as a function of genomic distance P_c_(*s*) and its derivative have been determined from the per-chromosome file generated by Hicstuff *pipeline*. The contact decay probability of the mitochondrial genome, the endogenous 2μ plasmid (as well as the artificial chromosomes in the case of ssHi-C libraries) were removed and the genome-average P_c_(s) and slope was computed using the *distance law* function of the Hicstuff package with default parameters within a reference window of 3 and 300 kb. Alternatively, the reference window was set to 3 and 100 kb when compared to ssDNA decay curves.

#### Quantification of DSB end-tethering

Sparse matrices in Graal or Cool format were load using the Hicstuff package *flexible_loader* function and converted in dense format using the *to_dense* function. The sum of contacts between two 60 kb windows (one on each DSB end and 70 kb away from *CEN5*) was computed. This sum was divided by the average of 11 equivalent control windows located at the same distance from their respective centromeres on chr. II, IV, VII, IX, X, XI, XII, XIII, XIV, XV and XVI. In the absence of DSB the ratio is ∼1. The Python code is available in the Jupyter notebook “Quantify left-right DSB ends chr5.ipynb”.

#### Quantification of coverage from Hi-C/ssHi-C reads

Quantification of coverage from *bam* alignment files generated with the Hicstuff *pipeline* function, which uses Bowtie2 in unpaired mate mode, were sorted and merged with Samtools *sort* and *merged* functions, respectively (Danecek *et al*, 2021). The coverage was computed in non-overlapping 500 bp bins using Tinycov *covplot* function (Matthey-Doret, 2025). The coverage was normalized onto the median genome coverage, and divided over that of samples lacking a DSB.

#### Generation of aggregated contact maps

Intra-chromosomal pile-ups between the top 500 cohesin-associated regions (CARs; detected from Scc1-V5 calibrated ChIP-seq from ref. (Piazza *et al*, 2021) using TinyMapper) were generated with Chromosight *quantify* with default parameters and a maximum distance of 50 kb (Matthey-Doret *et al*, 2020) from ICE-normalized Hi-C/ssHi-C matrices in cooler format binned at 1 kb and subsampled at 20 million contacts. Intra-chromosomal and inter-chromosomal centromere pile-ups were generated with Chromosight *quantify* centered on the 16 centromere positions or the intersection of centromere coordinates, respectively.

#### Gaussian contact probability model

Briefly, we modeled ssDNA as a homogeneous semi-flexible chain characterized by a persistence length *L*_*p*_ and a contour length *L*_*c*_. *L*_*p*_ corresponds to the typical lineic distance along the chain below which the ssDNA could be considered as rigid, and *L*_*c*_ to the lineic distance along the chain between the resection front and the DSB. Since the overall genome organization is not significantly affected during DNA repair compared to normal G2/M, we posited that the probability for a ssDNA probe to contact a dsDNA fragment depends on the relative 3D position of the resection front compared to the dsDNA locus in normal condition (no break) and on the probability of a site within the ssDNA emanating from the resection front to be found in close proximity to the dsDNA. The NPF thus plays the role of a fishing rod in the hands of a fisherman located at the resection front with a hook attached at the probe position. Assuming that the distribution of 3D relative positions between any pairs of loci to be Gaussian (Cattoni *et al*, 2017), we could then express the average contact probability between any ssDNA site with any dsDNA fragment as a function of the contact probability in normal conditions, of the statistics of position of the resection front and of the rigidity of the ssDNA (see **Supplementary Code 1**). The contact probability decay in normal conditions is given by Hi-C experiments. The statistics of resection front is extracted from a simple 1D stochastic model of the resection front around the DSB and fitted on the experimental Hi-C coverage data 2 and 4 hours post-DSB induction. The spatial properties of the NPF are given assuming a worm-like-chain statistics depending on *L*_*p*_ and *L*_*c*_. The only unknown parameter in the model is *L*_*p*_ that we then varied to fit the experimentally-measured fraction of *cis* contact by DSB-proximal and DSB-distal ssDNA sites. This departs from a previous inference using a similar model, in which *L*_*p*_ was varied to fit the experimentally measured fraction of *trans* contacts (Dumont *et al*, 2024). This modification allowed to overcome the altered distribution of *trans* contacts arising from the loss of end-tethering in the *srs2*Δ mutant. Both fitting methods give consistent results, yielding best-fit *L*_*p*_ = 20 nm in WT cells (Dumont *et al*, 2024). All the mathematical details of the model are provided in the **Supplementary Code 1**. The experimental data used to calibrate the model and determine the best fitting *L*_*p*_ values are available in **Dataset S2**.

#### Stochastic modeling of Rad51-ssDNA filament dynamics

A one-dimensional kinetic Monte-Carlo model called StochFilaments implemented in Python was developed to describe the dynamics of Rad51-ssDNA filament metabolism. The DNA is represented as a linear lattice of nucleotides, and Rad51 binding is modeled at the scale of a single Rad51 unit occupying 3 nucleotides (≈1.5 nm). Rad51 molecules organize into contiguous segments, corresponding to clusters of bound proteins. At each discrete and constant time step, three elementary processes are considered: nucleation, extension, and disruption. Nucleation occurs with probability *k*_*n*_ on any unoccupied site, creating a new segment of one Rad51 unit. Each existing segment elongates in the 3’ direction by exactly one Rad51 unit covering 3 nt per time step, until an occupied site or the end of the DNA lattice is reached. Disruption events occur with probability *k*_*d*_ per segment and are implemented in two alternative modes:

- *k*_*d1*_, a local dissociation where only the selected segment is removed.
- *k*_*d2*_, a directional dissociation where the selected segment and all its 5′ neighbors are removed, mimicking the processive action of a 3’-5’ helicase such as Srs2 (Rong & Klein, 1993; Kaniecki *et al*, 2017).

At each step the number of nucleation events is drawn from *Binomial(N_open, k*_*n*_) and the number of disruptions from *Binomial(N_seg, k*_*d*_), where *N_open* is the number of free sites and *N_seg* the number of existing segments. The two are applied in random order. Extension, by contrast, is deterministic, meaning that each eligible segment grows by one Rad51 unit per time step. The nucleation rate *k*_*n*_ and the dissociation rates *k*_*d1/2*_ are thus expressed relative to the growth rate. This approximation is justified experimentally, as nucleation rates are ∼10^-3^ that of the growth rate on protein-free ssDNA *in vitro* (Hilario et al, 2009; Candelli et al, 2014; Ma et al, 2017), a likely to be orders of magnitude lower on RPA-coated ssDNA. 40 independent replicates were simulated per parameter set. A default of 100,000 time-steps were simulated per replicate.

The model assumes that filament dynamics are dominated by local binding, polymerization, and dissociation rules, without explicit long-range interactions or three-dimensional conformational constraints. The ssDNA is treated as a one-dimensional substrate, each Rad51 segment is characterized by its length *I*_*i*_ and the total number of occupied sites *I* = ∑_*i*_*I*_*i*_ represents the ‘contour length’ of the full filament. For each simulation, the occupancy states of the filament were stored every 50 time-steps. From these occupancy matrices the number of Rad51 segments per molecule, the segment size distributions (mean, median, and maximum), the contour length, and the end-to-end distance were computed as time series for each cell and aggregated. End-to-end distances were computed, assuming that the full filament is a freely-jointed chain made by the segments and that each segment is rigid using:

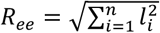

For each metric, the 40 replicates were aggregated and the steady-state regime identified by detecting the longest final portion of the trajectory over which the signal stayed flat (*i*.*e*. in which the fitted line changed by less than 2% of its mean). Runs that did not reach steady-state after 100,000 steps were extended 10- to 100-fold more time steps until a plateau could be reached.

Summary heatmaps were generated using the StochFilaments *postprocess* function.

#### Statistics & reproducibility

The statistical tests used are described in each figure legend. The statistical cut-off was set at 0.05. Statistical tests were performed with GraphPad Prism 10. No statistical method was used to predetermine sample size. No data were excluded from the analyses. The experiments were not randomized. The investigators were not blinded to allocation during experiments and outcome assessment.

### Data availability statement

Hi-C, ssHi-C and ChIP-seq sequencing reads and processed data as well as scripts and reference genomes used in this study will be made freely available at the time of publication.

#### Software availability

Other software used are available at:

- Hicstuff (version 3.2.4
- available at https://github.com/baudrly/hicstuff).
- ssHiCstuff (Mendiboure *et al*, 2026) (version 2.0.0 available at https://github.com/Piazzalab/sshicstuff).
- Parasplit (version 1.1.5 available at https://gitbio.ens-lyon.fr/LBMC/hub/parasplit/)
- Chromosight (Matthey-Doret *et al*, 2020) (version 1.6.3 available at https://github.com/koszullab/chromosight).
- Serpentine (Baudry *et al*, 2020) (version 0.1.3 available at https://github.com/koszullab/serpentine).
- Tinycov (version 0.3.1 available at https://github.com/cmdoret/tinycov).
- TinyMapper (version 0.10 available at https://github.com/js2264/tinyMapper).
- Integrative Genomics Viewer (Robinson *et al*, 2011) (version 2.8.3, available at https://igv.org).
- Bowtie2 (Langmead & Salzberg, 2012) (version 2.3.5.1 available online at http://bowtie-bio.sourceforge.net/bowtie2/).
- Samtools (Danecek *et al*, 2021) (version 1.3.1 available online at https://github.com/samtools/samtools).
- Cooler (Abdennur & Mirny, 2020) (version 0.9.1 available online at https://github.com/open2c/cooler).
- Pairtools (Open2C *et al*, 2024) (version 1.1.0 available online at https://github.com/open2c/pairtools).
- Deeptools (Ramírez *et al*, 2014) (version 3.5.5 available online at https://github.com/deeptools/deepTools).
- SciPy (Virtanen *et al*, 2020) (version 1.7.3 available online at https://github.com/scipy/scipy).
- Flowing Software (version 2.5.1 freely available online at https://flowingsoftware.com/download/).
- Graphpad Prism (version 10 commercially available at https://www.graphpad.com/).

## Acknowledgements

We thank members of the Piazza, Heyer, and Jost laboratories as well as Angela Taddei, Laurent Maloisel and Eric Coïc for helpful discussions. We are grateful to Boris Pfander for advices regarding the strand-specific ChIP-seq protocol. This research was supported by the European Research Council (ERC) under the European Union’s Horizon 2020 (ERC grant agreement 851006) to AP, the Agence Nationale de la Recherche to AP and DJ (ANR-23-CE12-0014-02), by the National Institutes of Health (R01GM58015, R35GM157976) to WDH, and by the France-Berkeley fund and the INSERM (IRP Filament) to AP and WDH.

## Author contributions

Conceptualization: AP; Experiments: AD, CD, JS, JL, AP; Data analysis: AP, JL; Modeling: NM, DJ; Data interpretation: NM, JL, WDH, DJ, AP; Supervision: WDH, AP; Funding Acquisition: DJ, WDH, AP; Manuscript - Writing: AP; Manuscript – Editing: DJ, WDH.

