## Supplementary figures for "Rad55-Rad57 and Srs2 regulate homology search onset, coordination, reach and inactivation"

### Mendiboure et al. Supplementary figures

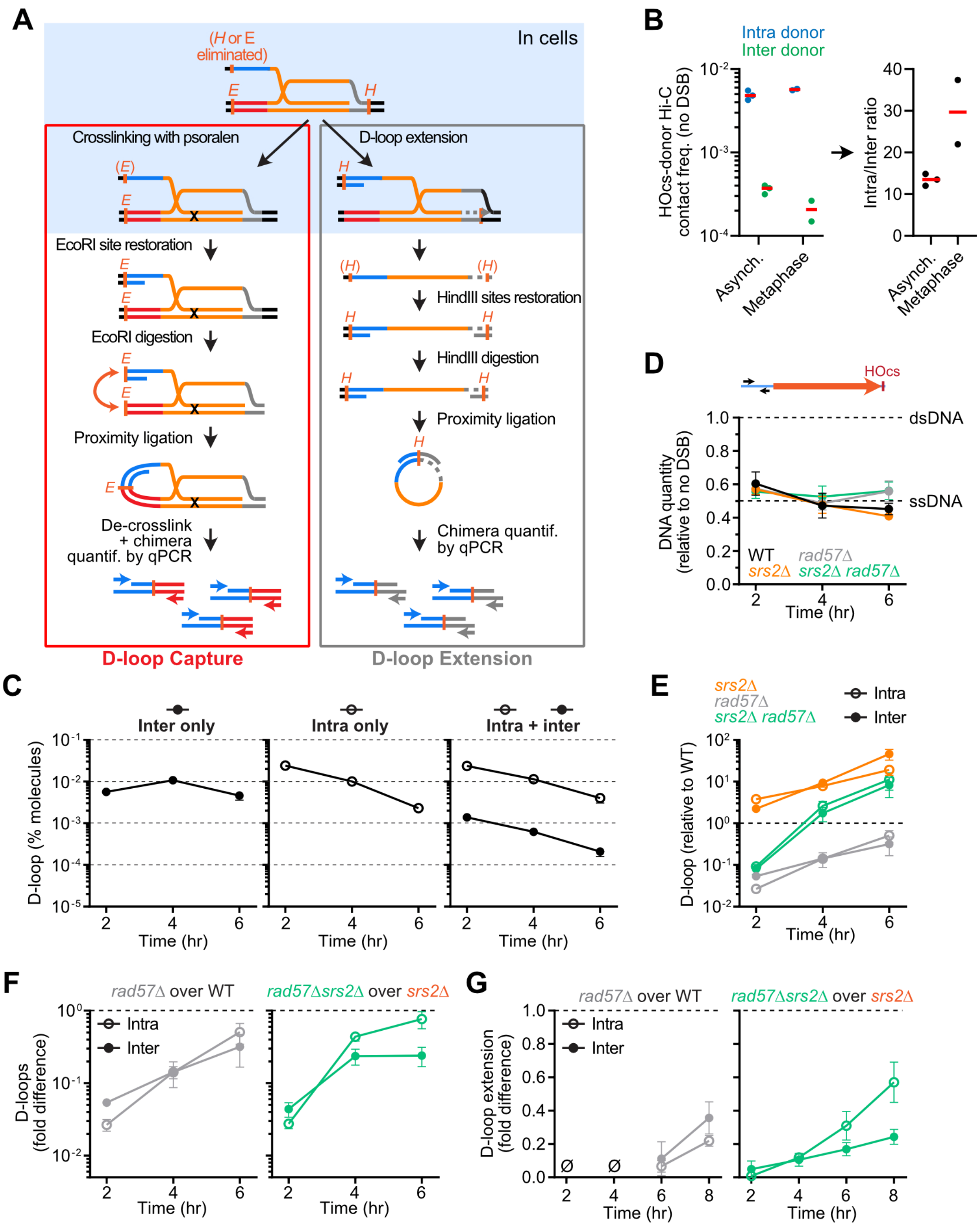

**Figure S1: DSB-donor spatial proximity partly alleviates the D-loop formation and extension defects of the *rad57Δ srs2Δ* mutant (related to Figure 1).**

A) Rationale of the D-loop Capture (DLC) and D-loop Extension (DLE) assays.
B) Hi-C contact frequency between the HOcs and the intra-chromosomal and inter-chromosomal donor loci in the absence of a DSB in asynchronous cells and in cells arrested at metaphase upon transcriptional repression of the *CDC20* gene. Data show individual biological replicates and median of Hi-C datasets published in (Dumont *et al*, 2024; Piveteau *et al*, 2026).
C) Absolute D-loop levels in WT strains bearing either or both an inter-chromosomal and an intra-chromosomal donor (APY266, APY826, and APY809). Data show mean ± SEM of n≥7 (inter donor only), n=1 (intra donor only) and n≥7 (intra+inter donor) biological replicates, and are replotted from data in (Djeghmoum & Piazza, 2025).
D) Quantification of ssDNA amounts over time in the same samples as in **Fig. 1B**. Data show mean ± SEM of at least n=3 biological replicates, except for *rad57Δ* at 2 hours (n=2).
E) Ratio of D-loop levels in mutants over WT strain determined in parallel, from data in **Fig. 1B**. Data show mean ± SEM.
F) Ratio of D-loop levels comparing *rad57Δ* over corresponding *RAD57* genotypes, calculated from data in **Fig. 1B**. Data show mean ± SEM.
G) Ratio of D-loop extension comparing *rad57Δ* over corresponding *RAD57* genotypes, calculated from data in **Fig. 1D**. Data show mean ± SEM. Ø: No DLE product detected.

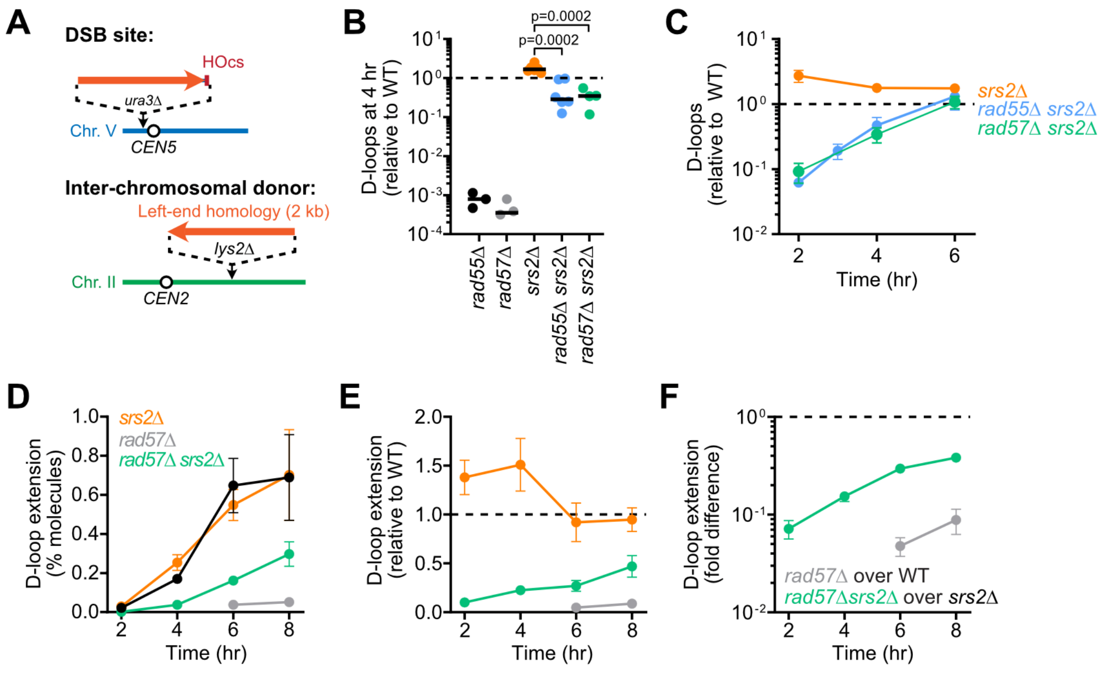


**Figure S2: D-loop formation and extension at an inter-chromosomal donor are delayed in the *rad55Δ srs2Δ* and *rad57Δ srs2Δ* mutants (related to Figure 1).**

A) Unrepairable site-specific DSB induction system with a 2 kb-long inter-chromosomal donor to the left DSB end.

B) D-loop-Capture levels 4 hours post-DSB induction in *rad55Δ* (APY11), *rad57Δ* (APY654), *srs2Δ* (APY773), *rad55Δ srs2Δ* (WDHY4609) and *rad57Δ srs2Δ* (APY1409) mutants relative to a WT strain (APY266/295) measured in parallel. Individual biological replicates and median are shown. Distributions were compared using a two-tailed unpaired Student t-test without Welch's correction.

C) D-loop-Capture levels are shown relative to a WT strain measured in parallel. Data show mean ± SEM of at least n=3 biological replicates.

D) D-loop extension levels at the intra- and inter-chromosomal donors in the same strains as in B). Data show mean ± SEM of at least n=3 biological replicates. No D-loop extension product was detected at 2 and 4 hours in the *rad57Δ* mutant.

E) Ratio of D-loop extension in mutants over the WT strain determined in parallel, calculated from data in D). Data show mean ± SEM.

F) Ratio of D-loop extension comparing *rad57Δ* over corresponding *RAD57* genotypes, calculated from data in D). Data show mean ± SEM.

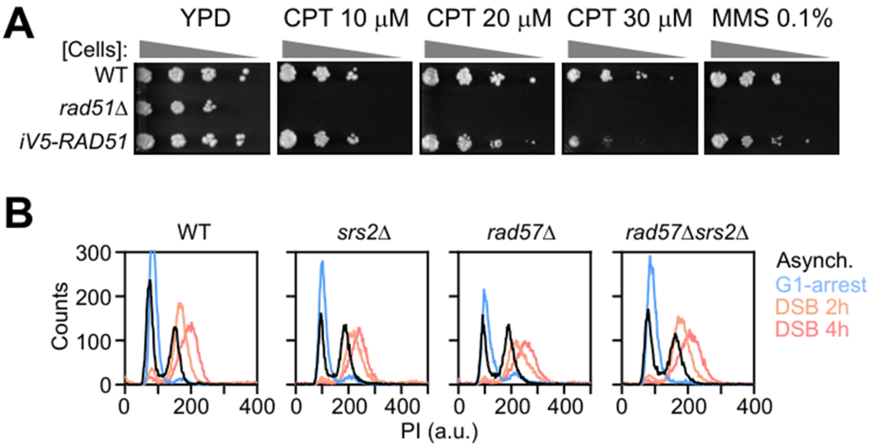


**Figure S3: The iV5-Rad51 construct is proficient for HR repair and DSB-induced G2/M-arrest (related to Figure 2).**

A) Spot viability assay of WT (APY266), *rad51Δ* (APY679) and *iV5-RAD51* (APY1895) strains in the presence of the genotoxic drugs camptothecin (CPT) and methyl methane sulfonate (MMS).

B) Validation of the cell synchronization in G1, S-phase release, and DSB-induced G2/M arrest by FACS of WT (APY1895), *srs2Δ* (APY2033), *rad57Δ* (APY2029), and *rad57Δ srs2Δ* (APY2037) strains expressing iV5-Rad51.

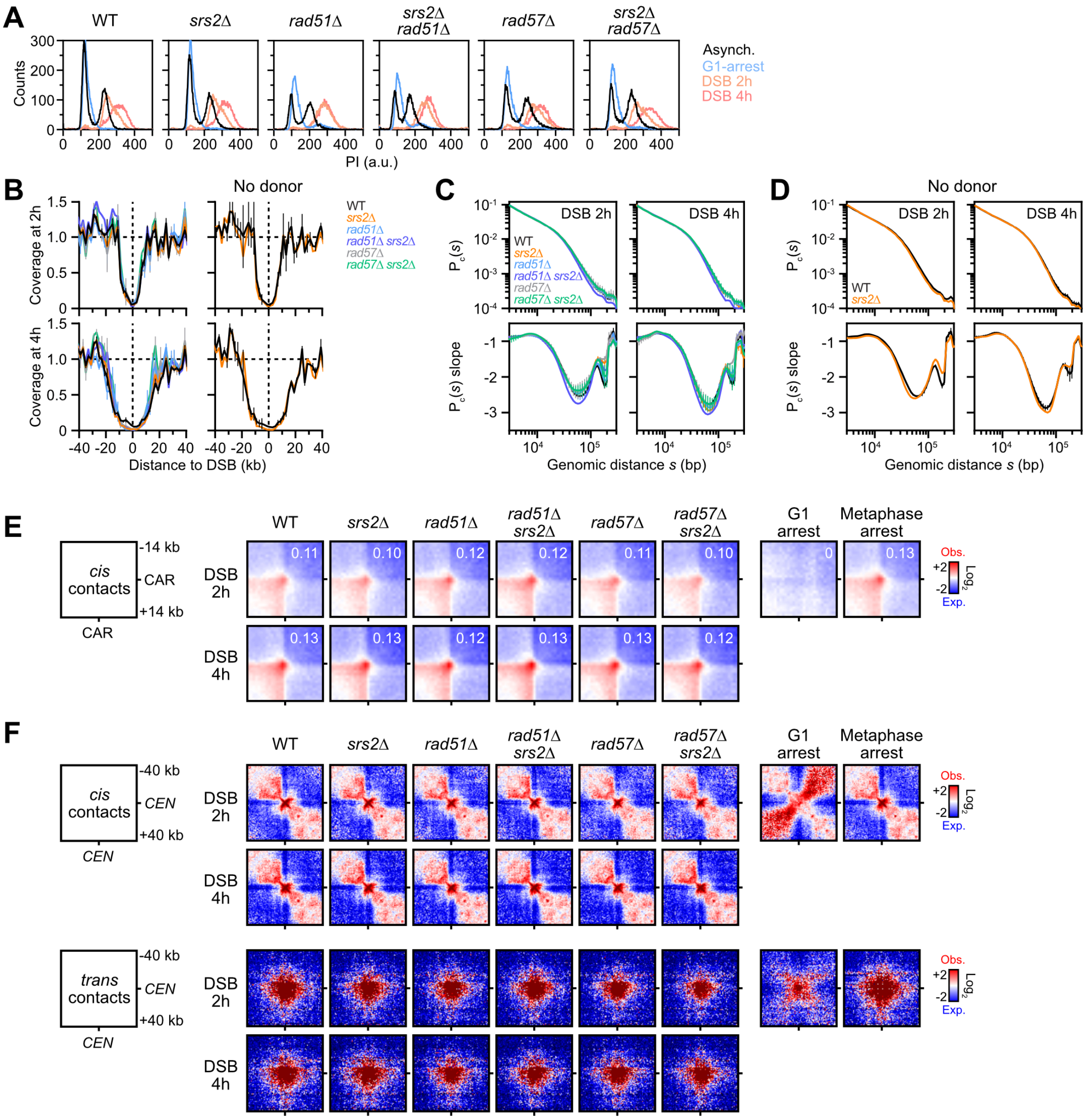
A
WT
srs2Δ
rad51Δ
srs2Δ rad51Δ
rad57Δ
srs2Δ rad57Δ
Counts
PI (a.u.)
Asynch.
G1-arrest
DSB 2h
DSB 4h
B
No donor
Coverage at 2h
Coverage at 4h
Distance to DSB (kb)
WT
srs2Δ
rad51Δ
rad51Δ srs2Δ
rad57Δ
rad57Δ srs2Δ
C
DSB 2h
DSB 4h
P_c(s)
P_c(s) slope
Genomic distance s (bp)
D
No donor
E
cis contacts
-14 kb
CAR
+14 kb
CAR
DSB 2h
DSB 4h
G1 arrest
Metaphase arrest
Obs.
Exp.
Log2
F
cis contacts
-40 kb
CEN
+40 kb
trans contacts
DSB 2h
DSB 4h
G1 arrest
Metaphase arrest

**Figure S4: Deletion of *RAD51*, *RAD57*, *SRS2*, or combination thereof does not affect resection, G2/M arrest, chromosome organization and cohesin-mediated chromatin loops (related to Figure 3).**

A) Validation of cell synchronization by FACS in WT (APY266), *srs2Δ* (APY773), *rad51Δ* (APY679), *rad51Δ srs2Δ* (APY1666), *rad57Δ* (APY654) and *rad57Δ srs2Δ* (APY1409) strains.
B) Hi-C coverage showing DNA resection tracts in strains in A) as well as WT (APY358) and *srs2Δ* (APY1644) strains lacking a donor. Data show mean ± SEM.
C) Genome-wide $P_c(s)$ and derivative of Hi-C contacts from strains in A-B). Data show mean ± SEM of n=2 biological replicates, except for WT (n=7) and *rad51Δ srs2Δ* (n=1).
D) Genome-wide $P_c(s)$ and derivative of Hi-C contacts from WT (APY358, n=2) and *srs2Δ* (APY1644, n=1) strains lacking a donor in B). Data show mean ± SEM.
E) Aggregated Hi-C contact maps between cohesin-associated regions (CARs) less than 50 kb apart, from strains in A). The average loop score is indicated.
F) Aggregated Hi-C contact maps of the 16 centromeric regions (top) and the inter-chromosomal *CEN-CEN* intervals (bottom), from strains in A).

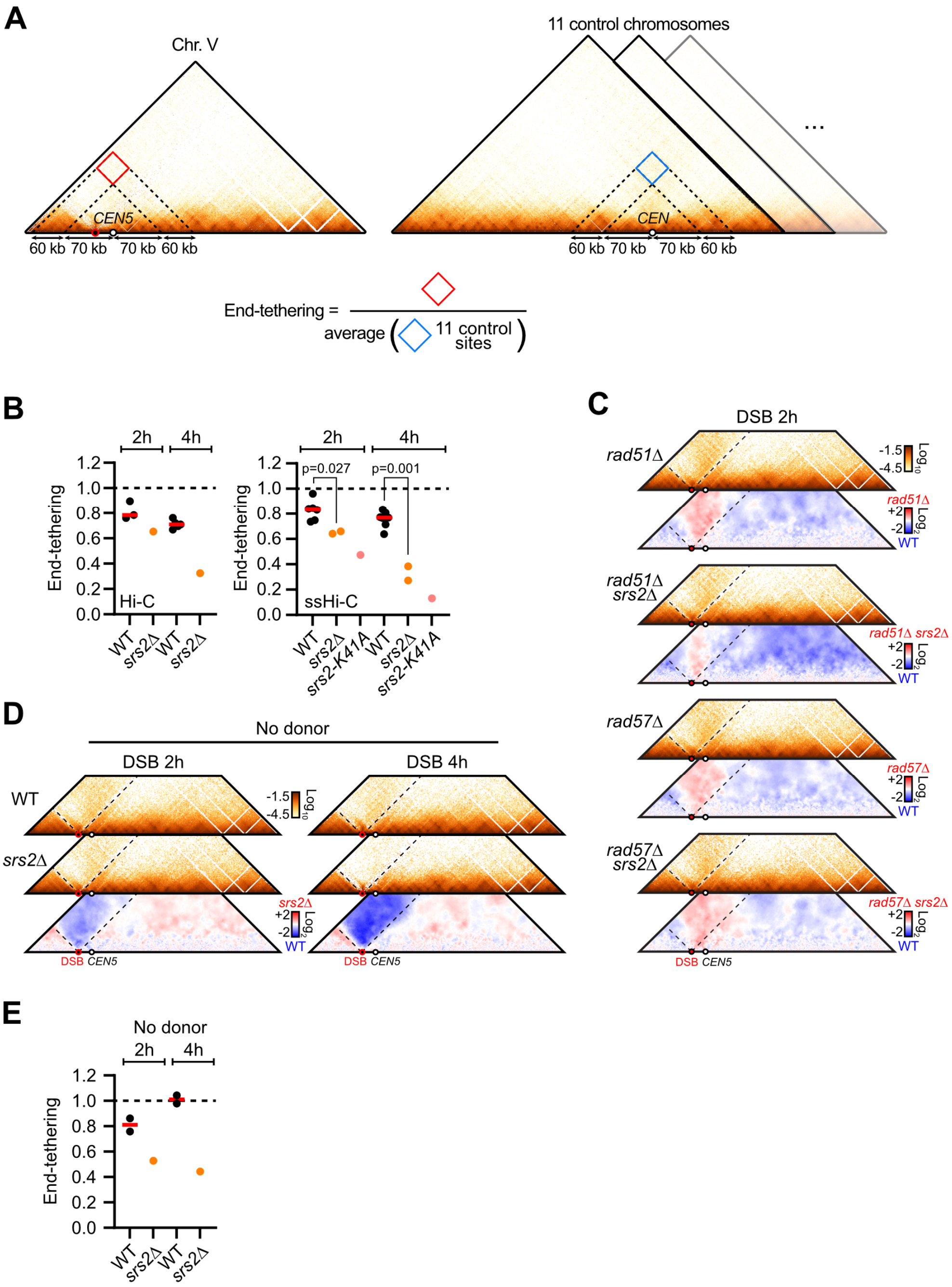

A
Chr. V
11 control chromosomes
CEN5
CEN
60 kb 70 kb 70 kb 60 kb
End-tethering =
average
11 control sites
B
2h 4h
End-tethering
Hi-C
ssHi-C
p=0.027
p=0.001
WT srs2Δ WT srs2Δ
WT srs2Δ srs2-K41A WT srs2Δ srs2-K41A
C
DSB 2h
rad51Δ
rad51Δ srs2Δ
rad57Δ
rad57Δ srs2Δ
WT
DSB CEN5
D
No donor
DSB 2h
DSB 4h
WT
srs2Δ
DSB CEN5
E
No donor
2h 4h
End-tethering
WT srs2Δ WT srs2Δ

**Figure S5: Loss of end-tethering in the absence of Srs2 depends on Rad51 and Rad55-Rad57, but not on the presence of a donor (related to Figure 3)**.

A) Rationale of DSB end-tethering quantification accounting for the brush-like organization of budding yeast chromosomes. The intersection of two 60 kb-long regions on each side of the DSB and at equidistance (70 kb) from *CEN5* is considered. The left interval is ~35 kb away from the DSB, *i.e.* out of the resection tract even 4 hours post-DSB induction (see **Fig. S4B**). The average of 11 control regions in the same configurations relative to their centromere is computed.
B) End-tethering from Hi-C and ssHi-C data in WT (APY266), *srs2Δ* (APY773) and *srs2-K41A* (WDHY4616) strains. Data show individual biological replicates and the median. Distributions were compared using an unpaired two-tailed Student t-test without Welch's correction.
C) Hi-C maps of chr. V in the *rad51Δ* (APY679, n=2), *rad51Δ srs2Δ* (APY1666, n=1), *rad57Δ* (APY654, n=2), *rad57Δ srs2Δ* (APY1409, n=2) mutants and corresponding ratio maps over a WT strain (APY266) at 2 hours post-DSB induction.
D) Hi-C maps of chr. V in WT (APY358, n=2) and *srs2Δ* (APY1644, n=1) strains lacking a donor, and corresponding ratio map.
E) Quantification of DSB end-tethering in strains in D). Data points show individual biological replicates and the median.

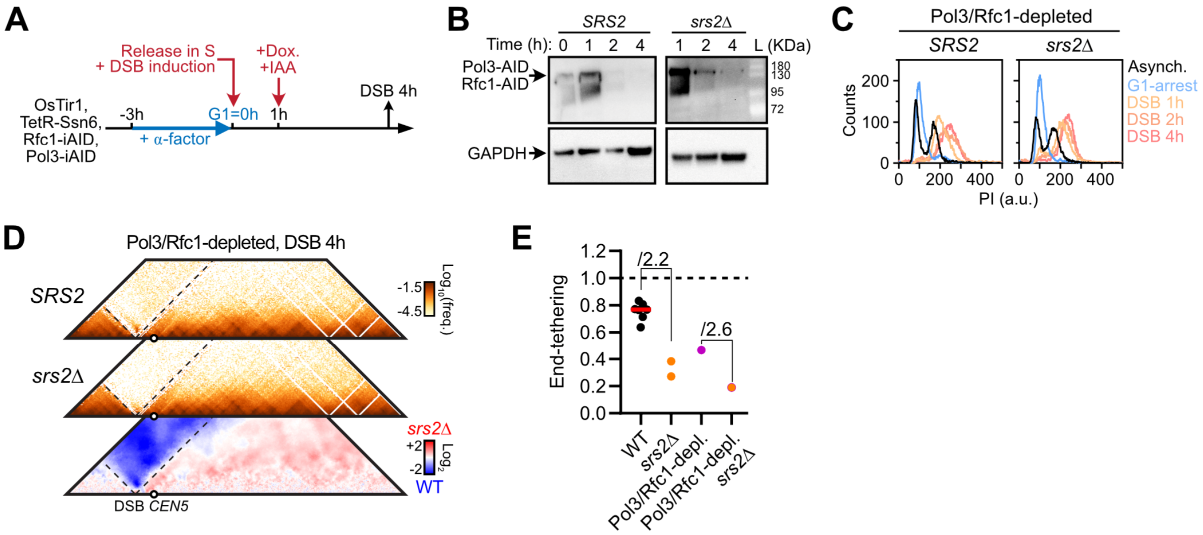
A
Release in S
+ DSB induction
+Dox.
+IAA
OsTir1,
TetR-Ssn6,
Rfc1-iAID,
Pol3-iAID
-3h
G1=0h
1h
DSB 4h
+ α-factor
B
SRS2
srs2Δ
Time (h): 0 1 2 4 1 2 4 L (KDa)
Pol3-AID
Rfc1-AID
180
130
95
72
GAPDH
C
Pol3/Rfc1-depleted
SRS2
srs2Δ
Counts
PI (a.u.)
Asynch.
G1-arrest
DSB 1h
DSB 2h
DSB 4h
D
Pol3/Rfc1-depleted, DSB 4h
SRS2
srs2Δ
Log10(freq.)
srs2Δ
Log2
WT
DSB CEN5
E
End-tethering
/2.2
/2.6
WT
srs2Δ
Pol3/Rfc1-depl.
Pol3/Rfc1-depl. srs2Δ

**Figure S6: Loss of end-tethering in the absence of Srs2 is independent of Pol3/Rfc1 (related to Figure 3).**

A) Synchronization, DSB induction and Pol3/Rfc1 depletion strategy.
B) Western blot validation of Pol3-iAID and Rfc1-iAID depletion in a WT (APY1350) and *srs2Δ* (APY1503) background. The western blots validating Pol3/Rfc1 depletion in a WT background are the same as in ref. (Dumont *et al*, 2024).
C) Validation of cell synchronization by FACS in Pol3/Rfc1-depleted WT and *srs2Δ* strains. The bulk of genomic DNA has been replicated when co-depletion of Pol3 and Rfc1 is induced.
D) Hi-C maps of chr. V in Pol3- and Rfc1-depleted WT (APY1350, n=1) and *srs2Δ* (APY1503, n=1) strains and corresponding ratio maps 4 hours post-DSB induction.
E) Quantification of end-tethering from Hi-C data in a WT strain (APY266), a *srs2Δ* mutant (APY773), a Pol3/Rfc1-depleted strain (APY1350) and a Pol3/Rfc1-depleted *srs2Δ* mutant (APY1503). Data show individual biological replicates and median.

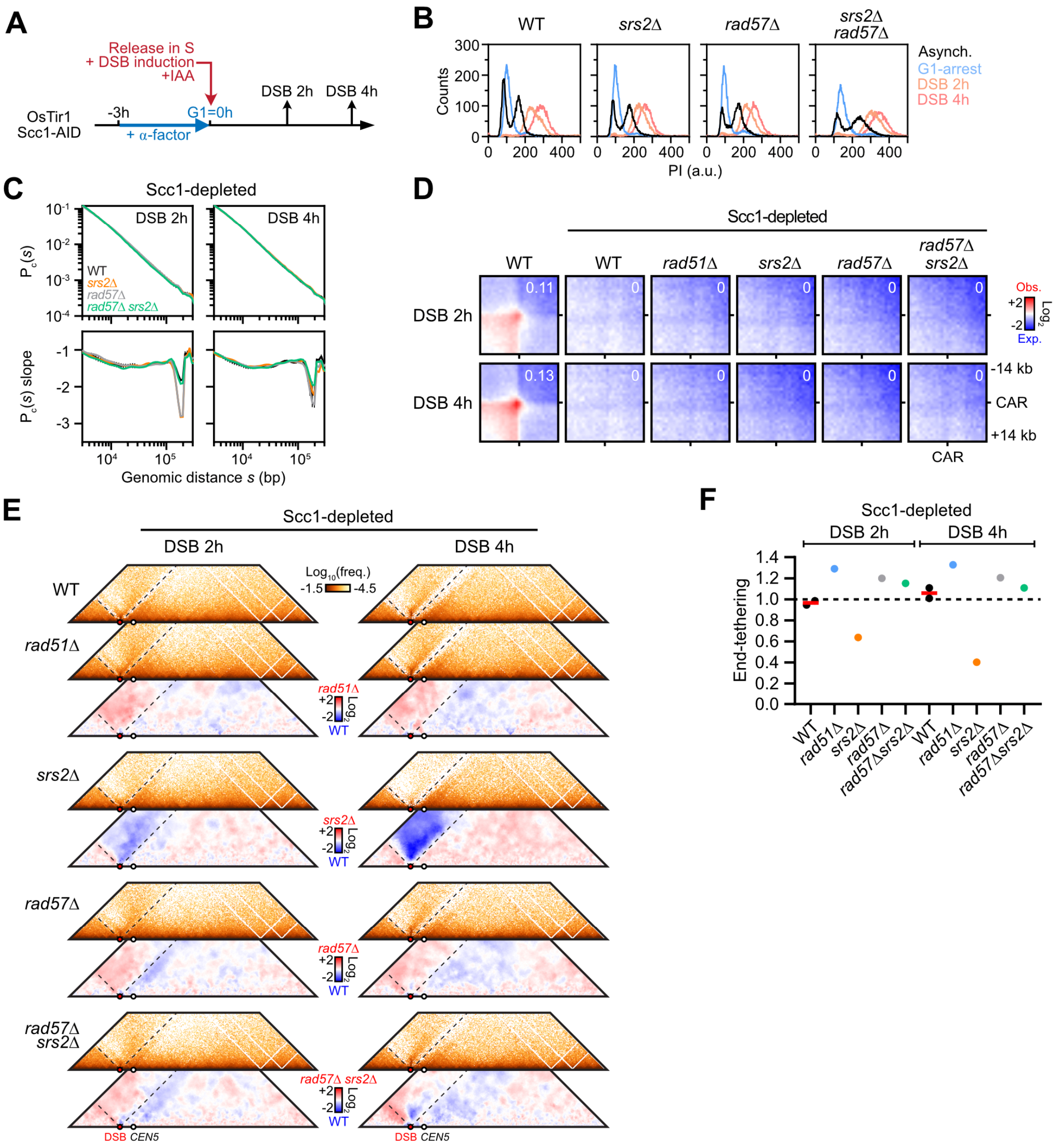
A
Release in S
+ DSB induction
+IAA
OsTir1
Scc1-AID
-3h
G1=0h
+ α-factor
DSB 2h
DSB 4h
B
WT
srs2Δ
rad57Δ
srs2Δ rad57Δ
Counts
PI (a.u.)
Asynch.
G1-arrest
DSB 2h
DSB 4h
C
Scc1-depleted
DSB 2h
DSB 4h
Pc(s)
WT
srs2Δ
rad57Δ
rad57Δ srs2Δ
Pc(s) slope
Genomic distance s (bp)
D
Scc1-depleted
WT
rad51Δ
rad57Δ srs2Δ
DSB 2h
DSB 4h
Obs.
Exp.
Log2
-14 kb
CAR
+14 kb
E
Scc1-depleted
Log10(freq.)
rad51Δ
srs2Δ
rad57Δ
rad57Δ srs2Δ
DSB
CEN5
F
End-tethering

**Figure S7: Loss of end-tethering in the absence of Srs2 is independent of Scc1 (related to Figure 3).**

A) Synchronization, DSB induction and Scc1 depletion strategy.
B) Validation of cell synchronization by FACS in Scc1-depleted WT (APY1481), *srs2Δ* (APY1467), *rad57Δ* (APY1479), and *rad57Δ srs2Δ* (APY1465) strains showing the maintenance of the DSB-induced G2/M arrest.
C) Genome-wide $P_c(s)$ and derivative of ssHi-C contacts in Scc1-depleted WT, *srs2Δ*, *rad57Δ*, and *rad57Δ srs2Δ* strains in E), showing a loss of typical cohesin-dependent enrichment of contacts in the 10-30 kb range (**Fig. S4C**) (Dauban *et al*, 2020).
D) Aggregated Hi-C contact maps between cohesin-associated regions (CARs) less than 50 kb apart showing the loss of loop signal in Scc1-depleted strains in E-F). The average loop score is indicated.
E) Hi-C maps of chr. V in Scc1-depleted WT (APY1481, n=2), *rad51Δ* (APY1500, n=1), *srs2Δ* (APY1467, n=1), *rad57Δ* (APY1479, n=1) and *rad57Δ srs2Δ* (APY1465, n=1) cells, from data subsampled to 16 million contacts and binned at 2 kb, and corresponding ratio maps of the mutants over the WT strain.
F) Quantification of end-tethering from ssHi-C data in Scc1-depleted WT (APY1481), *srs2Δ* (APY1467), *rad51Δ* (APY1500), *rad57Δ* (APY1479), and *rad57Δ srs2Δ* (APY1465) strains, from data in E). Data show individual biological replicates and median.

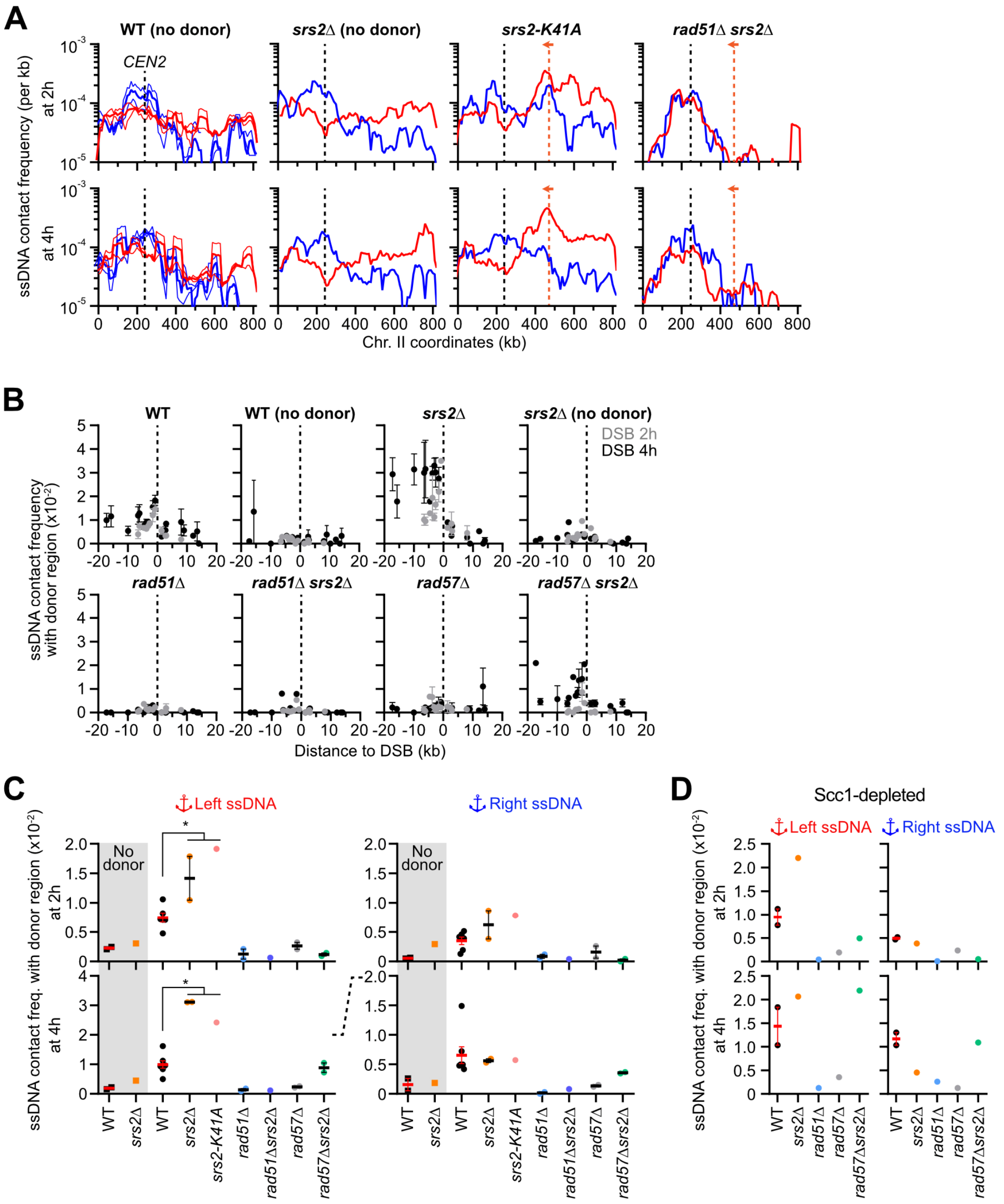
A
WT (no donor)
srs2Δ (no donor)
srs2-K41A
rad51Δ srs2Δ
CEN2
ssDNA contact frequency (per kb)
at 2h
at 4h
Chr. II coordinates (kb)
B
WT
WT (no donor)
srs2Δ
srs2Δ (no donor)
DSB 2h
DSB 4h
rad51Δ
rad51Δ srs2Δ
rad57Δ
rad57Δ srs2Δ
ssDNA contact frequency with donor region (x10⁻²)
Distance to DSB (kb)
C
Left ssDNA
Right ssDNA
No donor
ssDNA contact freq. with donor region (x10⁻²)
WT
srs2Δ
srs2-K41A
rad51Δ
rad51Δsrs2Δ
rad57Δ
rad57Δsrs2Δ
D
Scc1-depleted

**Figure S8: Srs2 promotes coordinated homology search by the left and right Rad51-ssDNA filaments (related to Figure 4).**

A) Distribution of left and right DSB-proximal ssDNA contact on chr. II in a WT strain and a *srs2Δ* mutant lacking a donor (APY358 and APY1644, respectively), and in a *srs2-K41A* and a *rad51Δ srs2Δ* mutant containing a donor (WDHY4616 and APY1666, respectively). Data show n=1 biological replicate, except for the WT strain lacking a donor (n=2). Data show mean (± SEM) of 10 kb bins smoothed over 5 bins.

B) Contact frequency of individual ssDNA sites with a 60 kb region surrounding the donor site on chr. II in a WT (APY266, n=6 or 7), a WT lacking a donor (APY358, n=2), a *srs2Δ* mutant (APY773, n=2), a *srs2Δ* mutant lacking a donor (APY1644, n=1), a *rad51Δ* (APY679, n=2), a *rad51Δ srs2Δ* (APY1666, n=1), a *rad57Δ* (APY654, n=2) and a *rad57Δ srs2Δ* (APY1409, n=2) mutant. Data show mean ± SEM.

C) Quantification of left and right DSB-proximal ssDNA contacts with the 60 kb region surrounding the donor site on chr. II, from ssHi-C data in B). Data show individual biological replicates and the mean ± SEM. * $p$-values < 0.05. Distributions were compared using an unpaired two-tailed Student t-test without Welch's correction.

D) Same is C), but from ssHi-C in Scc1-depleted strains. Data show individual biological replicates and the mean ± SEM.

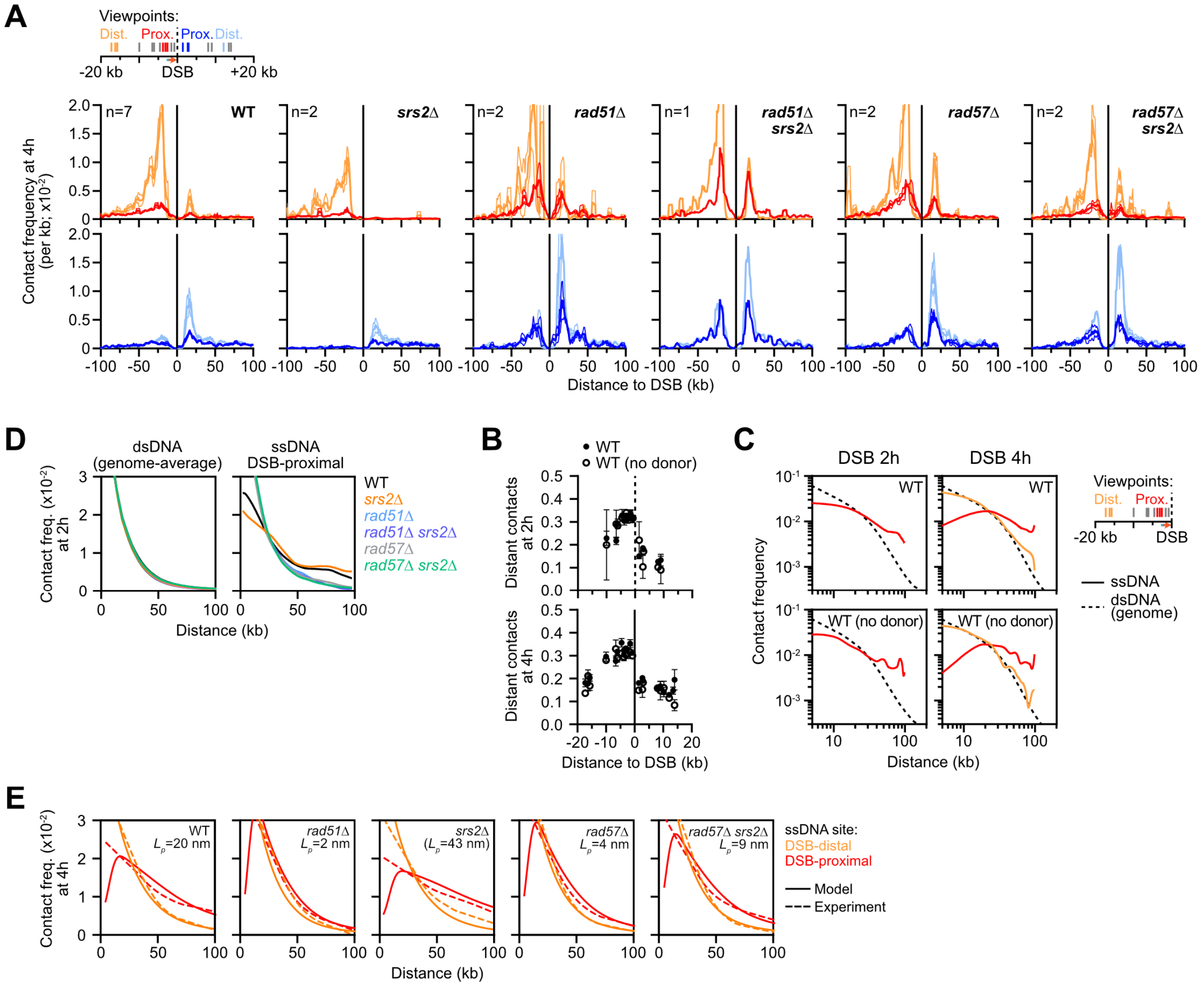
A
Viewpoints:
Dist.
Prox.
Prox.
Dist.
-20 kb
DSB
+20 kb
Contact frequency at 4h (per kb; x10^-2)
n=7
WT
n=2
srs2Δ
n=2
rad51Δ
n=1
rad51Δ srs2Δ
n=2
rad57Δ
n=2
rad57Δ srs2Δ
Distance to DSB (kb)
D
dsDNA (genome-average)
ssDNA DSB-proximal
Contact freq. (x10^-2) at 2h
WT
srs2Δ
rad51Δ
rad51Δ srs2Δ
rad57Δ
rad57Δ srs2Δ
Distance (kb)
B
WT
WT (no donor)
Distant contacts at 2h
Distant contacts at 4h
Distance to DSB (kb)
C
DSB 2h
DSB 4h
WT
WT (no donor)
Contact frequency
Distance (kb)
Viewpoints:
Dist.
Prox.
-20 kb
DSB
ssDNA
dsDNA (genome)
E
Contact freq. (x10^-2) at 4h
WT Lp=20 nm
rad51Δ Lp=2 nm
srs2Δ (Lp=43 nm)
rad57Δ Lp=4 nm
rad57Δ srs2Δ Lp=9 nm
ssDNA site:
DSB-distal
DSB-proximal
Model
Experiment
Distance (kb)

**Figure S9: Rad55-Rad57 and Srs2 together promote genome-wide homology search (related to Figure 5).**

A) Distribution of DSB-distal and DSB-proximal ssDNA contacts in the 200 kb region surrounding the DSB 4 hours post-DSB induction in WT (APY266), *srs2Δ* (APY773), *rad51Δ* (APY679), *rad51Δ srs2Δ* (APY1666), *rad57Δ* (APY654) and *rad57Δ srs2Δ* (APY1409) strains. Data show mean ± SEM of 1 kb bins smoothed over 5 bins. The number of biological replicates n is indicated. From data in **Fig. 5A, B**.

B) Contact frequency as a function of genomic distance for dsDNA genome-wide and DSB-proximal ssDNA on the left end side of the DSB in WT, *srs2Δ*, *rad51Δ*, *rad51Δ srs2Δ, rad57Δ* and *rad57Δ srs2Δ* strains 2 hours post-DSB induction, from data in **Fig. 5A, B**.

C) Proportion of ssHi-C contacts with distant genomic regions for individual ssDNA sites in a WT strain with (APY266, n=6 or 7) or without (APY358, n=2) a donor. Data show mean ± SEM.

D) LOWESS regression of the distribution of DSB-proximal and DSB-distal ssDNA contacts with chromatin in *cis* in a WT strain with (APY266) and without (APY358) a donor. The genome-wide $P_c(s)$ computed from dsDNA Hi-C contacts over the same interval is shown for comparison.

E) Comparison of experimental and best-fit modeling data of the distribution of DSB-proximal and DSB-distal ssDNA contacts with chromatin in *cis* 4 hours post-DSB induction. Data obtained with **Supplementary Code 1**. No $L_p$ could fit both the DSB-distal and DSB-proximal ssDNA contact distributions obtained in the *srs2Δ* mutant.

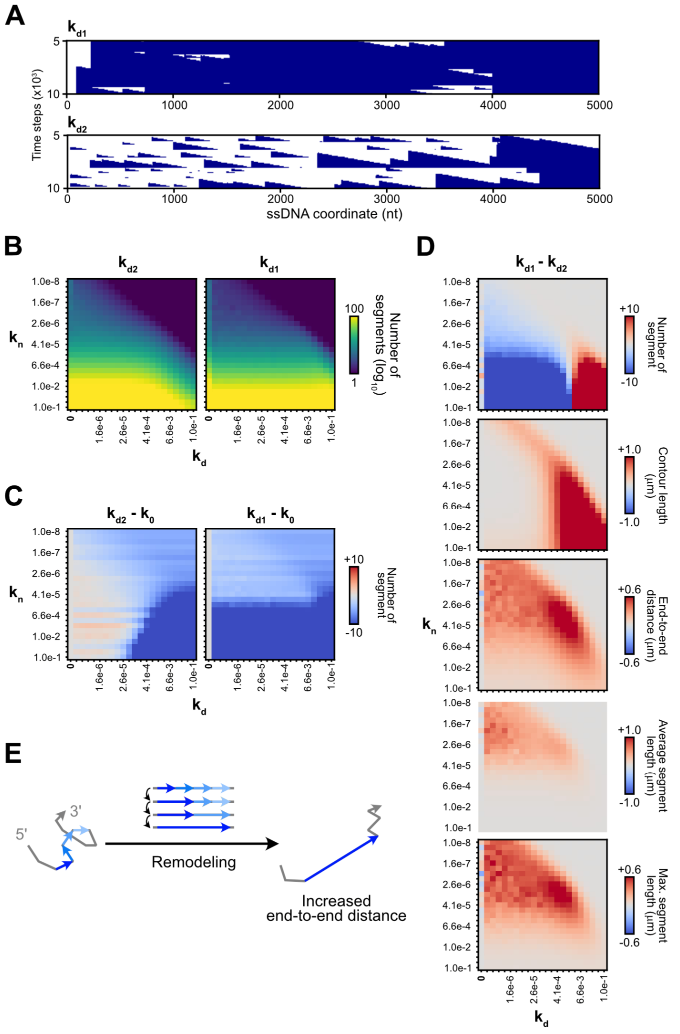


**Figure S10: Stochastic modelling of Rad51-ssDNA filament structural dynamics (related to Figure 7).**

A) Simulated kymographs of Rad51 filaments on 5 kb-long 5'-3' ssDNA molecules, corresponding to data in **Fig. 7B**. Blue marks occupancy by Rad51.

B) Heatmaps showing the steady-state number of Rad51 segments per ssDNA molecule with varying $k_n$, $k_{d1}$ and $k_{d2}$.

C) Effect of $k_{d1}$ and $k_{d2}$ (relative to no $k_d$) on the steady-state number of Rad51 segments per ssDNA molecule. From data in B).

D) Difference between $k_{d1}$ and $k_{d2}$. From data in B) and **Fig. 7C**.

E) Model for ssDNA stiffening upon reduction of Rad51 filament segmentation. Disruption of a Rad51 segment relieves a roadblock for the growth of the upstream segment. Iterative disruption-growth cycles stochastically lead to formation of a long Rad51-ssDNA segment, which increases the overall persistence length of the filament.
